# Contrasting impacts of urban and farmland cover on flying insect biomass

**DOI:** 10.1101/2020.09.16.299404

**Authors:** Cecilie S. Svenningsen, Diana E. Bowler, Susanne Hecker, Jesper Bladt, Volker Grescho, Nicole M. van Dam, Jens Dauber, David Eichenberg, Rasmus Ejrnæs, Camilla Fløjgaard, Mark Frenzel, Tobias Guldberg Frøslev, Anders Johannes Hansen, Jacob Heilmann-Clausen, Yuanyuan Huang, Jonas Colling Larsen, Juliana Menger, Nur Liyana Binti Mat Nayan, Lene Bruhn Pedersen, Anett Richter, Robert R. Dunn, Anders P. Tøttrup, Aletta Bonn

## Abstract

Recent studies report declines in biomass, abundance and diversity of terrestrial insect groups. While anthropogenic land use is one likely contributor to this decline, studies assessing land cover as a driver of insect dynamics are rare and mostly restricted in spatial scale and types of land cover. In this study, we used rooftop-mounted car nets in a citizen science project (‘InsectMobile’) to allow for large-scale geographic sampling of flying insects across Denmark and parts of Germany. Citizen scientists sampled insects along 278 10 km routes in urban, farmland and semi-natural (grassland, wetland and forest) landscapes in the summer of 2018. We assessed the importance of local to landscape-scale effects and land use intensity by relating insect biomass to land cover in buffers of 50, 250, 500 and 1000 m along the routes. We found a negative association of urban cover and a positive association of farmland on insect biomass at a landscape-scale (1000 m buffer) in both countries. In Denmark, we also found positive effects of all semi-natural land covers, i.e. grassland (largest at the landscape-scale, 1000 m), forests (largest at intermediate scales, 250 m), and wetlands (largest at the local-scale, 50 m). The negative association of insect biomass with urban land cover and positive association with farmland were not clearly modified by any variable associated with land use intensity. Our results show that land cover has an impact on flying insect biomass with the magnitude of this effect varying across spatial scales. Since we consistently found negative effects of urban land cover, our findings highlight the need for the conservation of semi-natural areas, such as wetlands, grasslands and forests, in Europe.

## Introduction

### Insect decline related to changes in land cover and land use intensity

Agricultural production and urbanisation have increased over the centuries, with at least three-quarters of the global land area currently affected by human activities (IPBES 2019). The IPBES Global assessment draws a sobering picture of global biodiversity decline associated with human activities; however, much of our understanding is based on vertebrates and plants. Yet, the majority of terrestrial animal species are insects (Stork, 2018). Changes in their biomass, abundance, and community composition could have diverse consequences, via alterations of food webs, nutrient recycling, pollination, and pest control, among others. Recent studies have found declining populations for terrestrial arthropod groups (e.g., Thomas *et al*. 2004; Hallmann *et al*. 2017, 2020; Valtonen *et al*. 2017; van Klink *et al*. 2020), with especially good evidence for some Lepidoptera, the order that is most commonly subject to the most long-term monitoring (e.g. Conrad *et al*. 2006; van Strien *et al*., 2019; Bell, Blumgart, & Shortall, 2020). However, at the same time, some insect taxa are expanding their distribution in Europe, including some dragonfly species, at least over recent decades (Termaat *et al*. 2019). Other insect groups, such as ants, seem to be able to persist even when exposed to extreme change (Guénard, Cardinal-De Casas & Dunn, 2015). However, we still lack a comprehensive view of how land cover and land use intensity affect insect populations (Seibold *et al.*, 2019). Few studies have simultaneously compared insect biomass across multiple different habitat types and at different spatial scales. Nonetheless, understanding relationships between insect biomass and land cover and land use is essential for conservation strategies aiming to mitigate insect loss.

### The effect of urbanisation on insect diversity and biomass

Arguably one of the most extreme land cover changes imposed by human activities is urbanisation (Seto, Güneralp & Hutyra, 2012). Across several insect taxa, Piano *et al.* (2020) found that urbanisation was associated with a decline in insect diversity at multiple spatial scales in Belgium. Similarly, a recent meta-analysis combining studies from across the world found a mean negative effect of urbanisation on terrestrial arthropod diversity and abundance, although the effect may differ among insect orders (Fenoglio, Rossetti & Videla, 2020). Long-term monitoring in Britain found lepidopteran biomass to be lower in urban sites compared to woodland and grassland (Macgregor *et al*., 2019). However, not all urbanisation is created equally. Cities with greater amounts of green space harboured higher insect pollinator abundances than cities with less green space (Turrini & Knop, 2015). Nor do all taxa respond in the same ways to urbanisation. For example, in one study, Hymenoptera showed higher species richness and flower visitation rates in urban areas compared to rural areas, with the opposite pattern exhibited by Lepidoptera and Diptera (Theodorou *et al*., 2020).

### Agricultural land use effects on insect diversity and biomass

Similar to urbanisation, land conversion for crop production has substantial consequences for biodiversity. In one study, Lepidopteran biomass was lower in arable areas compared to woodland and grassland sites (Macgregor *et al*., 2019). However, most studies on the impact of agricultural habitats have focused on comparisons among farming systems, e.g., conventional vs organic, rather than comparisons to semi-natural habitats (e.g. Kleijn & Sutherland, 2003; Bengtsson, Ahnström & Weibull, 2005; Bianchi, Booij & Tscharntke, 2006; Boutin, Martin & Baril, 2009). Overall, insect species richness has been reported as, on average, 30% higher in areas with organic farming, although the positive effects vary over spatial scales, and among taxa and functional groups (Bengtsson, Ahnström & Weibull, 2005). In Germany, Red List Lepidoptera biomass and species richness are twice as high in organic farmland compared to conventional farmland (Hausmann *et al*., 2020). Also, a high proportion of semi-natural areas within farmland, in essence, a more extensively used landscape, can have positive effects on insect taxa such as pollinators, that rely on the food and nesting resources such extensive landscapes provide (Cole et al., 2017). However, in general, it is less clear how much insect biomass in farmland (whether organic or conventional) differs from semi-natural areas, not to mention natural ecosystems, even though effects of agriculture or agricultural intensification are often invoked to explain insect declines (Goulson *et al.*, 2015; Goulson, Thompson & Croombs, 2018).

### Grassland, wetland, and forest land cover effect on insect diversity and biomass

Grassland, wetland and forests are often considered semi-natural because they are to some extent human-modified compared to natural ecosystems. In the cultural landscapes of Europe, all these habitats have variable land use histories resulting in a continuum from semi-natural to highly managed. Most forests in Western Europe are often managed for timber production (McGrath *et al*., 2015) and grasslands are often improved for extensive agri-environmental practices, i.e. managed for high biomass yields of high energy content fodder. These land covers might be expected to have greater abundance and biomass of insects than urban and agricultural areas. Indeed, sites with more forest cover showed weaker temporal declines of insect biomass in Germany (Hallman et al., 2017); while Lepidoptera biomass in woodland sites across Britain increased over a period of 10 years (Macgregor et al., 2019). Still, some studies show no significant differences in insect abundances and diversity between managed and semi-natural forests (Young & Armstrong, 1994; Watt, Barbour & McBeath, 1997; Humphrey *et al.*, 1999), although natural habitat structures related to deadwood, veteran trees and glades have been shown to be crucial for threatened specialist species (Heilmann-Clausen & Christensen, 2004; Lassauce, *et al*., 2011).

### Study approach and expectations

Drivers of fluctuations in insect populations are challenging to assess since long-term spatio-temporal population data are rare (De Palma *et al*., 2018). However, analysis of spatial patterns might indicate which land cover and land use changes are most harmful to insects. In this study, we investigate drivers of insect biomass by examining spatial patterns across two European countries, Denmark (northern Europe) and Germany (middle Europe). Denmark is covered by 74% highly human-modified landscapes with 61% agricultural areas, 13% settlements and infrastructure, and the remaining landscape mainly covered by 13% forests and 11% semi-natural areas (Statistics Denmark, 2019). Germany is similarly covered by highly human-modified landscapes, with over 50% of the land area used for farming, and the remaining area primarily used for forestry (31%) and human settlements and infrastructure (13.7%) (German Federal Statistics Office, 2015). For our study, we motivated citizen scientists to sample flying insects with car-nets as part of the InsectMobile project. Car nets have been employed for biting flies, mosquito and beetle sampling by professionals and amateurs for more than half a century (e.g. Bidlingmayer, 1966; Dyce, Standfast and Kay, 1972; Roberts and Irving-Bell, 1985), but have not been used as a standardised insect sampling method before. Our approach has the advantage of allowing multiple land covers to be sampled nearly simultaneously across large scales in a uniform and standardised way.

To our knowledge, this is the first study to simultaneously assess the effects of multiple land covers and land use intensities on insect biomass. We compared insect biomass among major land cover types: urban, farmland, grassland, wetland and forest across Denmark and parts of Germany. We focused on insect biomass for several reasons: it aligns with reported declines of insect biomass (Hallman *et al.*, 2017); it is a relevant measure for ecosystem functioning (Barnes *et al.*, 2016), and it is a measure of resource availability for higher trophic levels. Overall, we hypothesised that insect biomass would be lower in areas with more human-modified land cover and more intense land use. Specifically, we assume (H1) lower biomass in farmland areas compared to open semi-natural habitats (wetland and grassland) due to agricultural practices such as pesticide use, homogenisation, and ploughing and harvesting, either directly killing insects or removing habitats and resources for insects. Further, we assume (H2) that urban cover would have the lowest insect biomass among all land covers due to the high proportion of impervious surfaces and the low proportion of blue and green space, meaning limited food, nesting and breeding resources. Finally, we assume (H3) that insect biomass within highly-modified land cover types would be negatively associated with increasing land use intensity, reflected by variables such as intensity of farming and urban structural composition, i.e. larger cities and urban green space.

## Materials & Methods

### Citizen science sampling with car nets

Flying insects were sampled by standardised nets attached to the rooftop of cars. The car net is funnel-shaped with a detachable sampling bag at the far end for sample collection. Metal guy line adjusters enable adjustment to car length and allow the net to be used on most car types (Figure 1).

**Figure 1:**
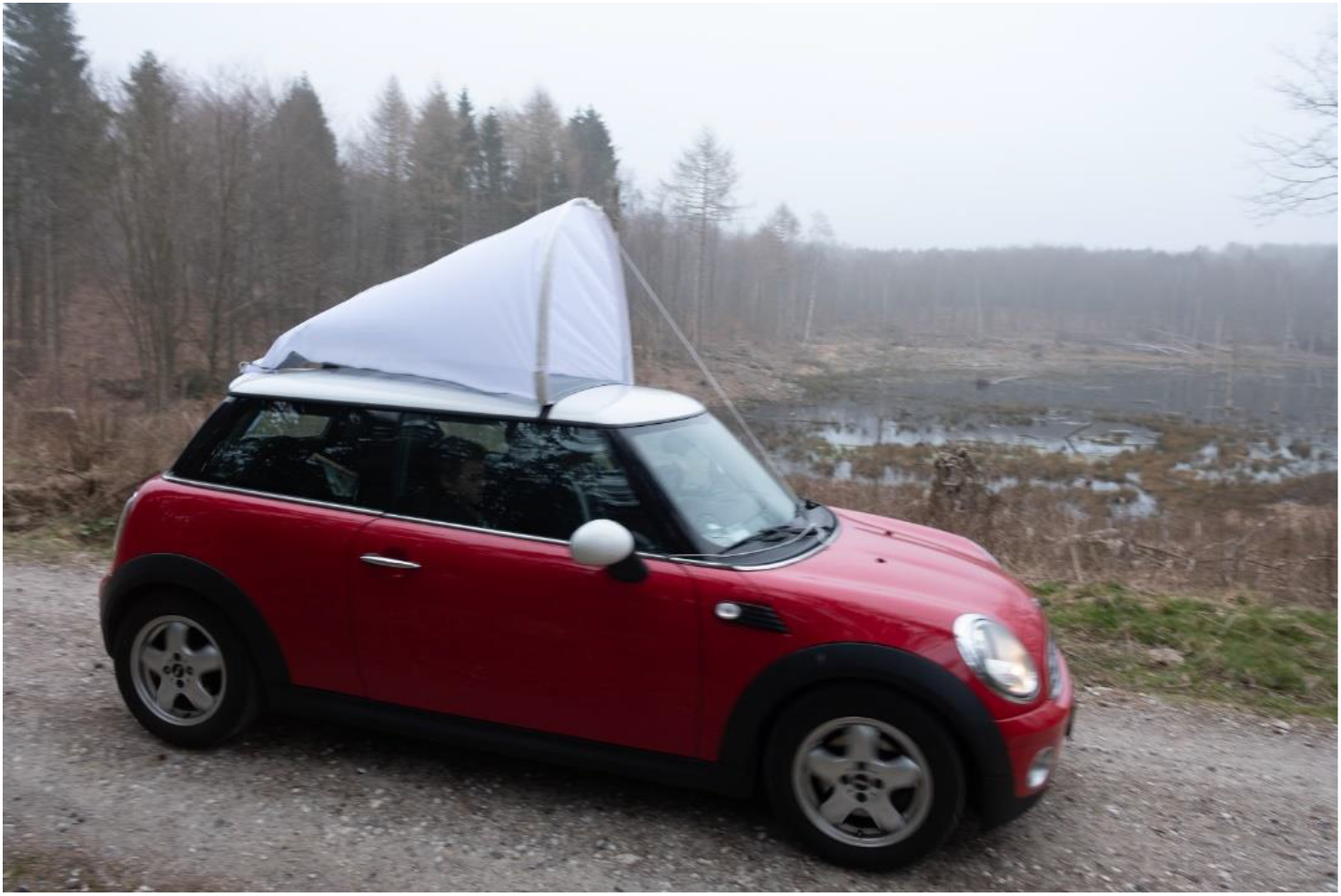
Car net used to sample flying insects. Picture from The Natural History Museum of Denmark’s promotional video. Photo: Anders Drud | Natural History Museum of Denmark.

Citizen scientists were recruited by the Natural History Museum of Denmark in Denmark (NHMD) and for a scoping study by the German Centre for Integrative Biodiversity Research (iDiv) in Germany during spring 2018. The citizen scientists received a simple sampling protocol as well as video tutorials and FAQ sheets along with the sampling equipment (Supplementary Information (SI) V).

Sampling was carried out by 151 Danish and 29 German citizen scientists along 211 routes from 1 - 30 June 2018 in Denmark, and along 67 routes between 25 June - 8 July 2018 in Germany (Figure 2). Sampling of each route was carried out in two time intervals during the day: between 12-15 h (midday) and between 17-20 h (evening) with a maximum speed of 50 km/h and weather conditions of at least 15°C, an average wind speed of maximum 6 m/s and no rain. Samples were placed in 96% pure ethanol and sent back to NHMD and iDiv by the citizen scientists.

**Figure 2:**
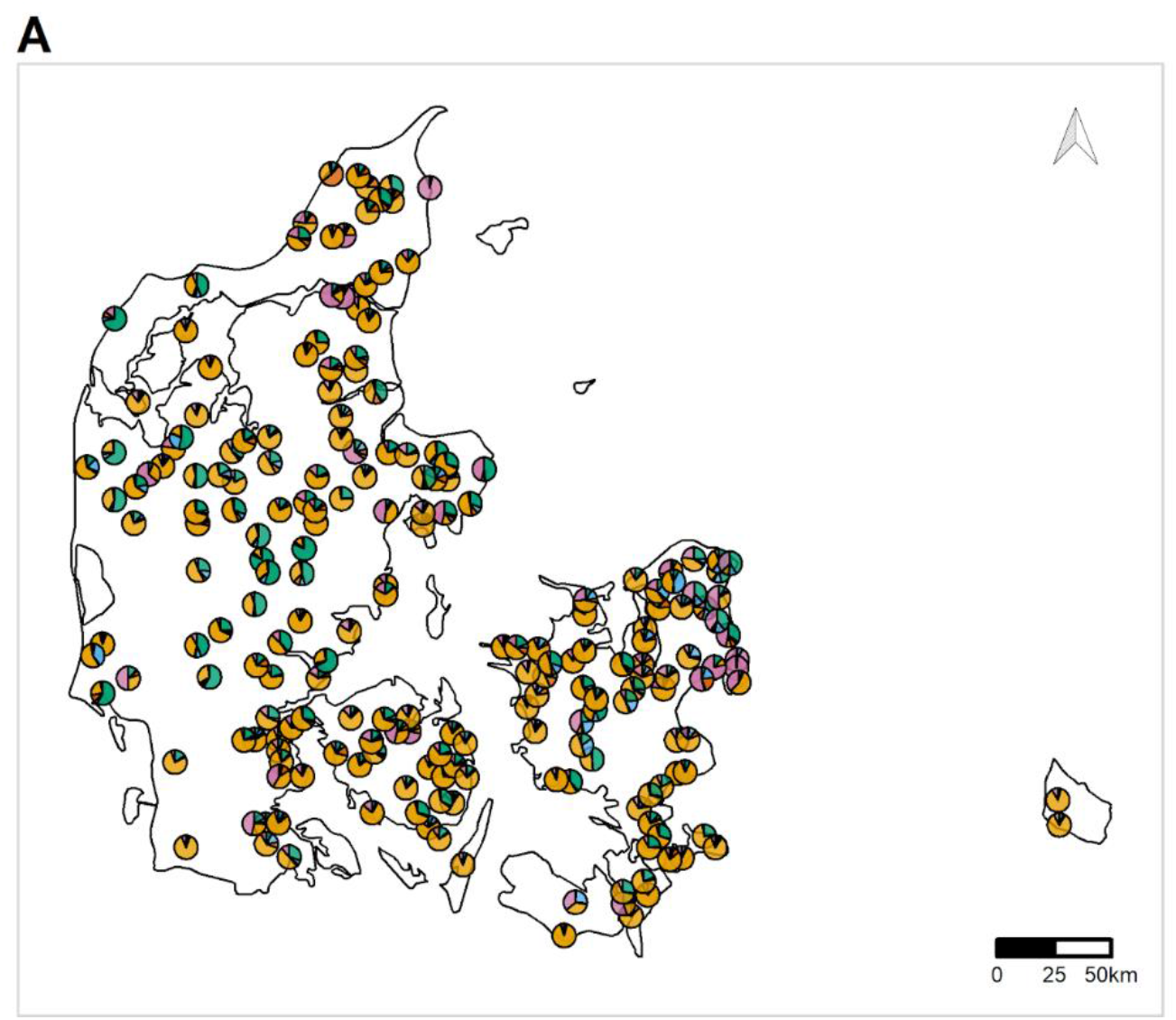
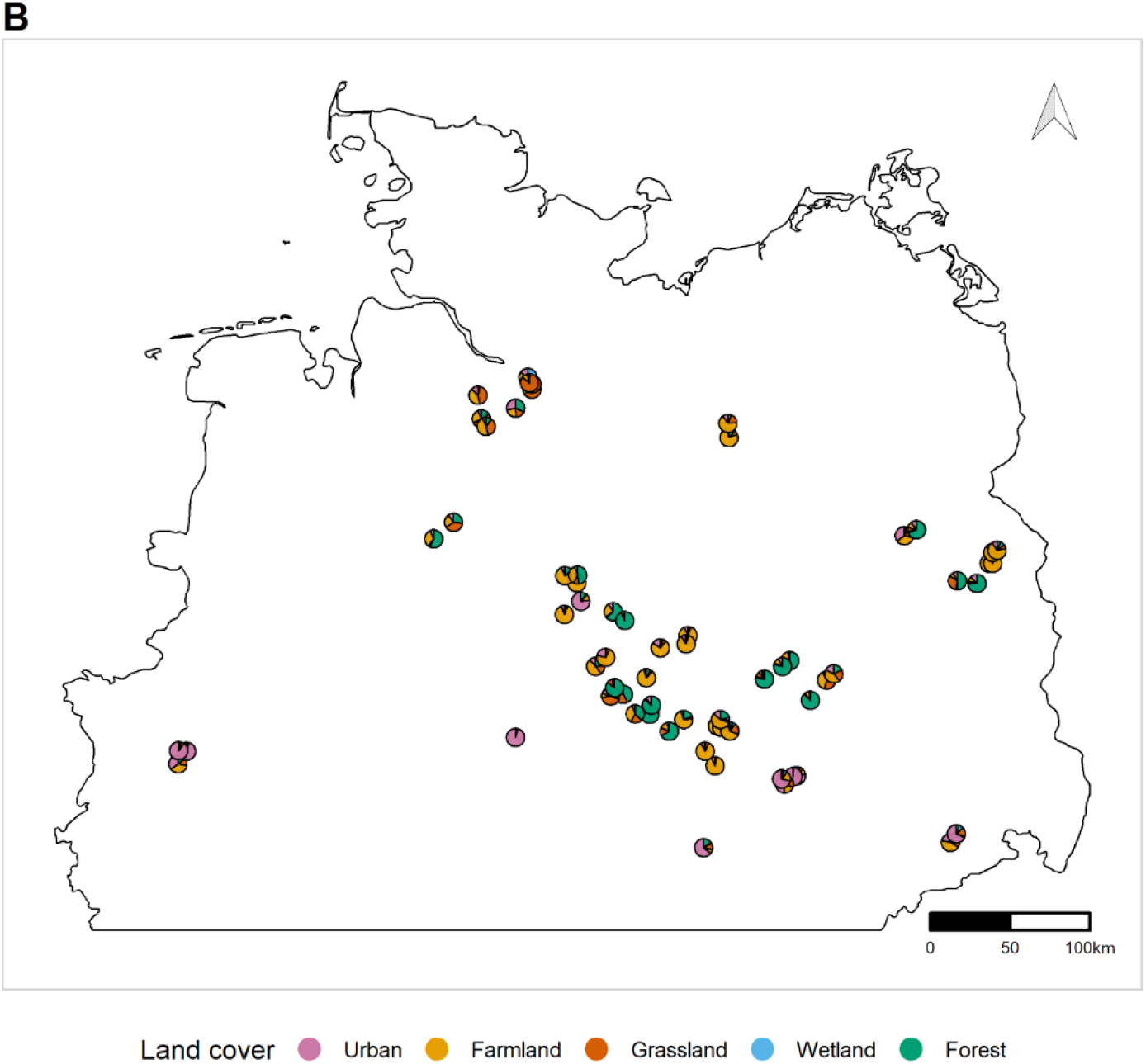
Location of car net sampling routes in two European countries A) Denmark (211 routes) and B) Germany (67 routes). Pie chart points illustrate the proportional land cover at the 1000 m buffer for each sampling location.

### Route design

Across both countries, routes were designed with a length of 5 km across five land cover types: farmland, grassland, wetland, forest and urban areas. Each sampling event covered 10 km length in total - either driven in one direction or 5 km driven in both directions to cover the total length. The routes were constructed in ArcGIS and QGIS using information from Google Earth, Google Maps, OpenStreetMap (OSM), including data from Danish authorities on land cover types in Denmark, and also using the German ATKIS data (Amtliches Topographisch-Kartographisches Informationssystem) in Germany. The different land cover data sources were used to assess the land cover along the routes to ensure as much homogeneity in the chosen land cover type as possible. Routes were adjusted, if needed, following incorporation of local area knowledge of the citizen scientists about land use, accessibility, road condition and safety. In a few cases in Germany, routes were shorter due to topographical limitations (e.g., extent of wetland and urban areas) and were therefore driven several times back and forth to achieve a total length of 10 km.

### Dry weight of bulk insect samples

In the laboratories of the NHMD and iDiv, insects were removed from the sampling bag with a squeeze bottle containing 96% EtOH and forceps. Empty 15 or 50 ml centrifuge tubes were weighed, and the insects were transferred to the tubes. The insects were dried overnight at 50 °C in an oven (>18hrs), and the tubes containing the dry insects were weighed to obtain the sample biomass (in mg).

### Environmental data

According to Seibold *et al.* (2019), the effect of land cover levels off at a 1000 m buffer for grassland and forest sites, we, therefore, extracted land use predictors for insect biomass from four buffer zones for each route: 50 m, 250 m, 500 m, and 1000 m in five categories; urban, farmland, grassland, wetland, and forest. Land use predictors were compiled into land cover categories. A comprehensive overview of land cover categories and their definitions are listed in Supplementary Information I. Land cover classifications were aligned across the Danish and German data to the same categories.

Land use intensity data for Denmark were extracted for farmland and urban routes. The farmland category consisted of crop types compiled into three overall categories: extensive, semi-intensive, intensive, and agricultural areas with no associated crop type. Extensive farmland is, e.g. fallow land etc., semi-intensive farmland is, e.g. orchards etc., and intensive farmland is, e.g. wheat, rye, beans, etc. Grass leys (rotational grassland in an agricultural area to ensure soil fertility) were included in all three intensity categories, whereas semi-natural grassland only consisted of grasslands under the Danish Protection of Nature Act Section 3. The three overall farmland categories in Denmark were further compiled into organic and conventional farming practices. For the available German data, it was not possible to make the distinction between grass leys and semi-natural grassland or farmland practices. The urban category for Denmark consisted of various building type categories, such as multistory buildings, residential areas, commercial areas and inner-city areas. Both multistory buildings and inner-city cover are only found for larger cities. These data were not available for Germany.

We extracted potential stop variables to account for sampling heterogeneity introduced by the number of stops along each route. We obtained the number of traffic lights or stops of any type (e.g. roundabouts, pedestrian crossings, stop signs, railroad crossings) within a 25-30 m buffer using OSM. For Danish routes, we obtained the number of roundabouts using data from the Danish Map Supply provided by SDFE (Agency for Data Supply and Efficiency) (GeoDenmark-data), since data on roundabouts in Denmark was limited to three records in OSM.

Mean hourly temperature and wind was extracted for each route including date and time band from the nearest weather station using the rdwd R package for German routes. For Danish routes, temperature, average wind speed, and sampling time were registered by the citizen scientists.

### Statistical analyses

The German and the Danish datasets were analysed separately while applying the same modelling approaches and methods to enable comparison.

#### Correlation and PCA

We first investigated the correlations among the land cover variables to explore land use gradients and assess whether multicollinearity would be an issue in multiple regression models. We investigated this by calculating pairwise Pearson correlation coefficient as well as principal components analysis (PCA). Correlations among land cover types were strongest between urban and farmland, with increasing farmland associated with decreasing urban cover (Denmark, r= −0.6; Germany, r= −0.46, both calculated for the 1000 m buffer). However, since the correlations among the land cover types were not strong (see SI IV: Figure 4.1), and hence none were redundant, we analysed the land cover variables as separate variables, but later considered the patterns with the land covers simplified to the first two PCA axes (see SI II). We used a varimax-rotated PCA to maximise the variation explained by each axis, using the psych R package (Revelle, 2020). Using the same model structure as below (equation 1), we used the first two PCA axes, as described above, as land cover explanatory variables in an additional set of models (SI II). Results from correlation tests and PCA can be found in the supplementary information.

#### General model

To test the impact of land cover on insect biomass, we analysed log biomass (+1, since there were a few zeros) as the response in mixed-effects models assuming a normal distribution, with land cover or land use variables as our main explanatory variables. To control for other factors causing variation in insect biomass, we included the day of the year, time band (midday vs evening), time of day (centred around each time band, and then nested within time-band as a predictor), weather variables (temperature and wind) and other measures of possible sampling variation (log-transformed number of traffic lights, or other stops) (hereafter, called controlling variables). Additionally, to account for potential non-independence of data points, we included random effects for route and citizen scientists (i.e., driver and car). The mixed-effects models were fit using lmer in the lme4 R package (Bates *et al.*, 2015).

Hence, the general form of the mixed-effects model was:

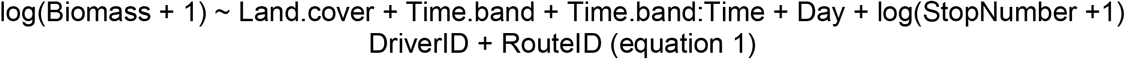

We consistently found no effect of weather variables (probably because of little variation, as the samples were taken under similar weather conditions), and therefore they were not included in the final models.

#### Spatial autocorrelation

Since the sampling points were spatially-structured, we investigated whether the models of insect biomass should account for spatial autocorrelation. We plotted correlograms and tested for spatial autocorrelation with Moran’s I (simulated residuals from the lmer model, DHARMa R package (Hartig, 2020)) but did not find evidence for spatial autocorrelation in the residuals of the fitted model of equation 1 (p = 0.3). Moreover, we also used a generalised least squares model (GLS, in R package nlme (Pinheiro *et al.*, 2020)), with the same response and explanatory variables described above and the geographic coordinates of each route as an exponential spatial correlation structure (nugget = TRUE). These models produced very similar results and models without the spatial term had a lower AIC. Based on these findings, we analysed the findings of the model without the explicit spatial structure.

#### Land cover as ecological predictors

Using models of the general form of equation 1, we tested the effect of each land cover variable. We used % coverage of each land use type (see documentation in SI I) within the four different buffer zones (50 m, 250 m, 500 m, and 1000 m) around each route, representing local to landscape-scale effects. To facilitate comparison of the effects of each land cover within and across countries, covariates were kept in their original units; hence, effect sizes of the land cover relate to change in biomass per 1% land cover change. Because some of the variables were skewed, we also checked the effects of applying square-root transformations to the land cover data.

#### Simple regression models

We first tested the effect of each land cover and buffer combination (5 land covers x 4 buffer widths) on insect biomass in simple regression models (i.e., one land cover variable per model, but including controlling variables of time, day and stops as well) (SI II: Figure 2.1). We used these simple models to identify the best buffer width (i.e., one with the largest effect size) for each land cover.

For the Danish data, we found a grassland outlier route containing around 40% grassland cover, where all other routes with grassland contained less than half of that cover (<20%). We excluded this route from the analysis, as it could introduce bias in our models (see SI II for model outputs and visualisation with the outlier).

#### Multiple regression models

##### Full model

We then built a linear mixed-effects model that included all five of the land cover variables (at the best buffer width for each one) and the controlling variables day of the year, time band, time of day, and log-transformed number of traffic lights or stops. We examined variation inflation factors to check whether collinearity among explanatory variables (i.e., variable redundancy) was an issue.

##### Best fit model

We identified the best fit model using AIC, i.e. the model with the lowest AIC, and ran the analysis with the modified models for each country (see included variables in both the full and the best fit model in Table 1). To examine the partial effects of each land cover variable, we used the effects R package (Fox & Weisberg, 2018) to predict the change in biomass with increased land cover for each land cover type, controlling for effects of other land covers as well as controlling variables, at their mean values (Figure 4), based on the model output from the full model.

**Table 1:** Regression coefficients of the linear mixed-effects model on insect log(biomass+1) (significant variables in bold). The full model includes all land cover and controlling variables. The best fit model was identified by the lowest AIC. All land cover variables were kept in their original units to facilitate interpretation. Shown is the mean (standard error) of each regression coefficient. Explained variation by the fixed effects in each model indicated in percent.

| Variable | Denmark (full model: 37%) <sup>▽</sup> | Denmark (best fit model: 33%) | Germany (full model: 36%) <sup>▽</sup> | Germany (best fit model: 30%) |
| --- | --- | --- | --- | --- |
| Urban - 1000 m | -0.013 (0.009) | **-0.017 (0.007) \*** | -0.028 (0.024) | **-0.040 (0.008) \*** |
| Farmland - 1000 m | **0.012 (0.005) \*** | **0.014 (0.005) \*** | 0.024 (0.016) | - |
| Grassland - 1000 m | **0.054 (0.019) \*** | **0.058 (0.018) \*** | 0.018 (0.021) | - |
| Wetland; DE: 1000 m, DK: 50 m | 0.051 (0.029) <sup>Δ</sup> | **0.057 (0.028) \*** | 0.005 (0.044) | - |
| Forest: DE: 250 m, DK: 250 m | **0.013 (0.085) \*** | **0.016 (0.005) \*** | 0.014 (0.013) | - |
| Day of year | **0.028 (0.006) \*** | **0.027 (0.006) \*** | -0.046 (0.042) | - |
| Time band: midday vs evening | **0.33 (0.09) \*** | **0.32 (0.09) \*** | **0.383 (0.115) \*** | **0.416 (0.116) \*** |
| Time within band (change in biomass per minute within time band) | Midday: -0.0005 (0.002)<br>**Evening: 0.006 (0.001) \*** | - | Midday: 0.0002 (0.002)<br>**Evening: 0.005 (0.002) \*** | - |
| Number of Stops/Traffic lights | -0.26 (0.19) | - | 0.4013(0.3217) | - |

\* $p < 0.05$ , <sup>Δ</sup> $p < 0.1$ , <sup>▽</sup> Generalised variance inflation factor for the full models; DE: 5 for grassland and stops, 10 for farmland, urban and forest, DK: 2.87 for stops and forest, 5.4 for urban cover, 4.8 for farmland, 1 for wetland, 1.2 for grassland.

We additionally tested whether the effect of land use depends on time band (i.e., a time band:land use interaction) as different insect communities are expected to be sampled at different times of the day, which may respond differently to land cover.

#### Land use intensity as ecological predictors

For routes dominated by urban or farmland, we further investigated whether variables associated with the intensity of land use within the 1000 m buffer explained variation in insect biomass. We restricted this analysis to the Danish routes because of the larger sample size. We first selected the routes where the dominant land cover (i.e., the highest proportion among land covers) was farmland within the 1000 m buffer for farmland intensity analysis, or urban within the 1000 m buffer, for urban intensity analysis. In this subset, 34 routes were dominated by urban areas, and 255 routes were dominated by farmland. We tested for correlation between land use variables by calculating pairwise Pearson correlation coefficient and PCA (see SI III).

To account for the association between general land cover and the land use intensity variables, we calculated the proportional cover of the land use intensity variable within the land cover variable (i.e., the proportion of green space within the urban land cover.) We then constructed models similar to equation 1, but included an interaction term between the overall land cover and the proportional land use intensity variables (equation 2). These interactions tested whether the effect of urban cover depended on the land use intensity properties of the urban cover, and similarly whether the effect of farmland cover depends on the land use intensity properties of the farmland cover:

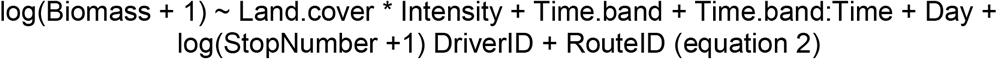

We examined whether there was a strong correlation between land covers and land use intensity variables after calculating the proportional cover of the intensity variables (SI III: Figure 3.4 & 3.8) and removed highly correlated variables in urban land use intensity analysis from the model.

**Figure 3:**
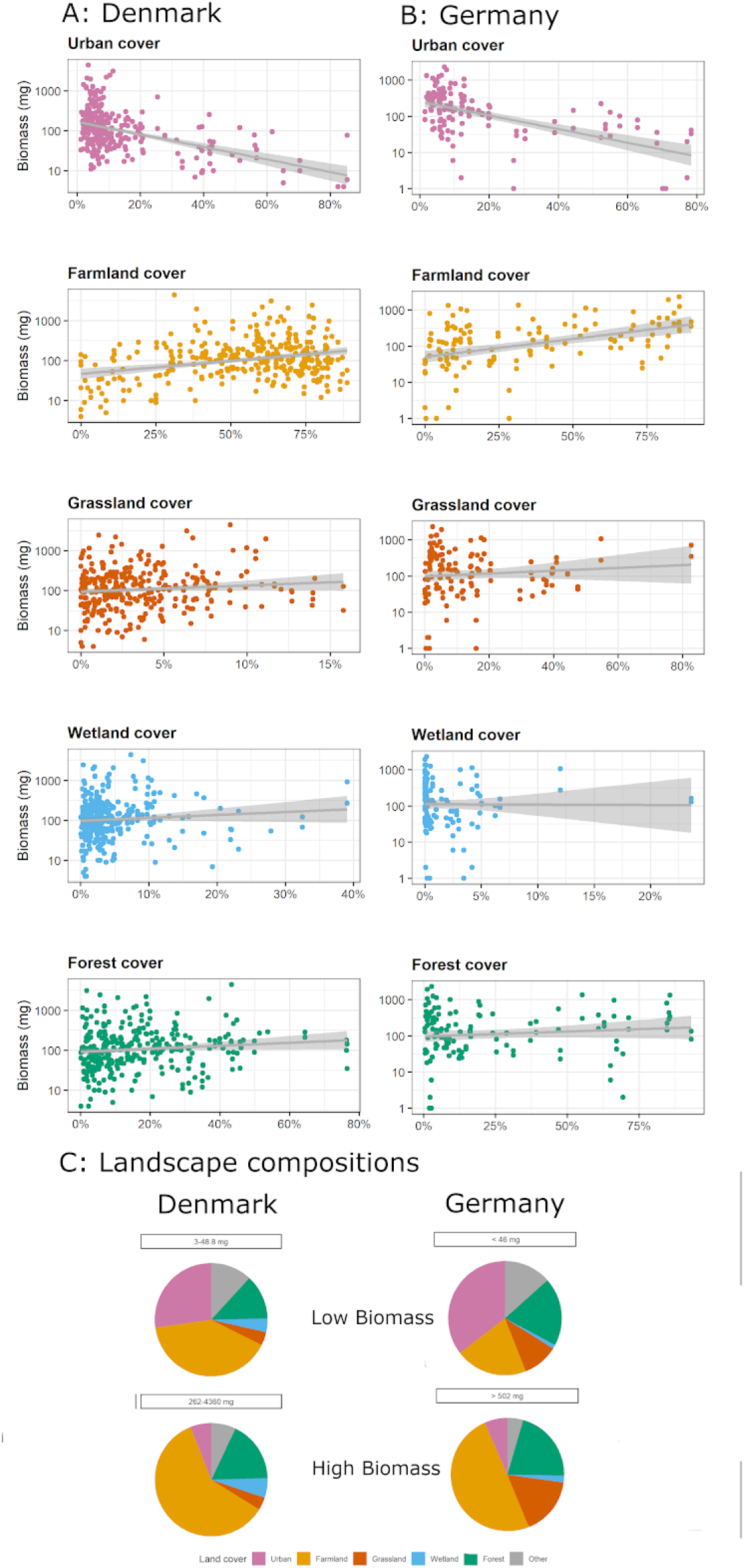
Scatterplots show the simple relationships between percent of each land cover and insect biomass. A) Denmark, B) Germany. C) Pie Charts show the mean land cover composition of routes with the lowest 20% quantile and upper 20% quantile of biomass samples.

All analyses were carried out in R (version 3.6.3).

## Results

### Land cover

We found a negative effect of urban land cover on insect biomass and higher biomass in all rural land covers, including farmland cover. Especially at the broader landscape scale, we found significant associations.

In Denmark, largest effect sizes for urban, farmland and grassland were associated with buffers of 1000 m as well as for 250 m for forest and 50 m for wetland (SI II: Figure 2.1A). In Germany, all land covers except forest had largest effect sizes associated with 1000 m buffers; forest cover had similar effect sizes with buffers between 250, 500 and 1000 m (SI II: Figure 2.1B). The dominant land cover types within the routes were farmland (mean coverage of 54% in Denmark, and 37% in Germany), urban (mean coverage of 12% in Denmark and 21% in Germany) and forest (mean coverage of 16% in Denmark and 26% in Germany), which reflect the coverage of these cover types in the two countries.

### Denmark

In the best fit model, we found a positive effect of wetland, grassland, forest and farmland on insect biomass, and a negative effect of urban land cover on insect biomass. Furthermore, we found a positive effect of increasing biomass throughout the month of June and higher biomass in the evening compared to midday. Fixed effects of land cover type and control variables explained 33% of the variation in biomass. Results were similar when land cover types with skewed distribution were transformed by a square-root transformation. The mean landscape composition for samples with high biomass (within top 20% of biomass samples, >262 mg) was dominated by farmland cover. In comparison, the mean landscape composition for samples with low biomass (within the bottom 20% of biomass samples, <48.8 mg) was dominated by urban areas as well as farmland (see Figure 3C).

In the full model, we found positive effects of farmland, forest and grassland cover on insect biomass, and a trend towards a positive effect of wetland on insect biomass (Table 1 & Figure 4A). The negative effect of urbanisation was, however, not significant. The fixed effects explained 37% of the variation in the model. In addition, we found a positive effect of sampling day with an increase in biomass throughout June, higher biomass in the evening compared to midday and an increase in biomass within the three-hour evening sampling (Table 1). The urban cover had a high correlation with potential stops along the routes (SI IV: Figure 4.1).

**Figure 4:**
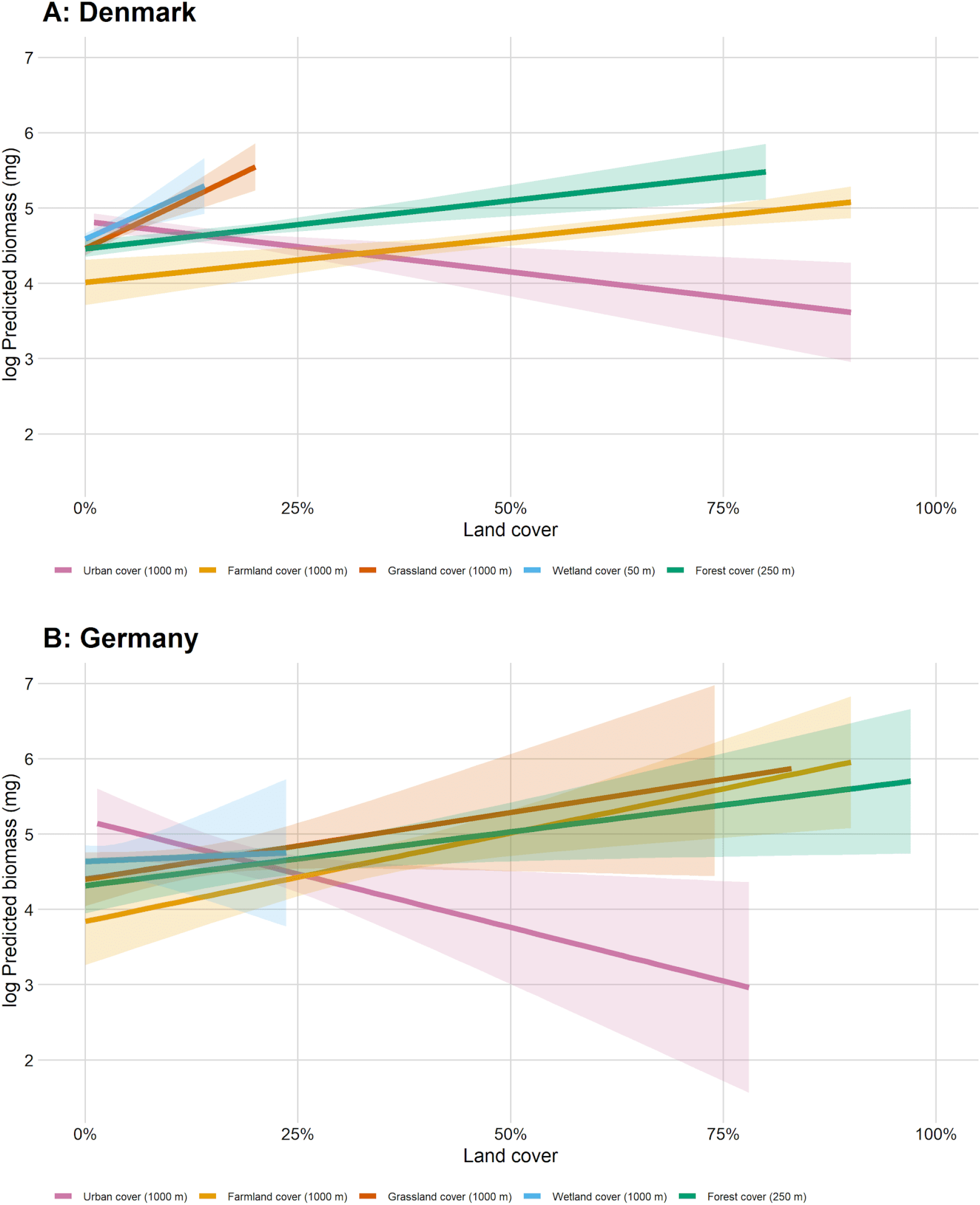
Partial effects of each land cover when all other predictors are held fixed at their means for (A) Denmark and (B) Germany. Predicted log(biomass+1) (mg) on the y-axis and proportional land cover on x-axis. Based on the full model for each country to illustrate the relative effect of each land cover. Shaded areas around each line is the standard error for the fit.

In the composite land cover analysis, the two axes of the varimax rotated PCA were driven by an urbanisation gradient (axis 1) and a forest gradient (axis 2) (SI II: Figure 2.2). We found a significant negative effect of the urbanisation gradient on insect biomass (p = 0.002), but no effect of the forest gradient (p = 0.31) (SI II: Table 2.1). The fixed effects in this model explained a third (34%) of the variation in insect biomass among routes.

Since we found an effect of timeband (more insects in the evening; Table 1, Figure 5A), we explored whether the effect of land cover differed with sampling time, but we did not find any evidence of an interaction between land cover and timeband.

**Figure 5:**
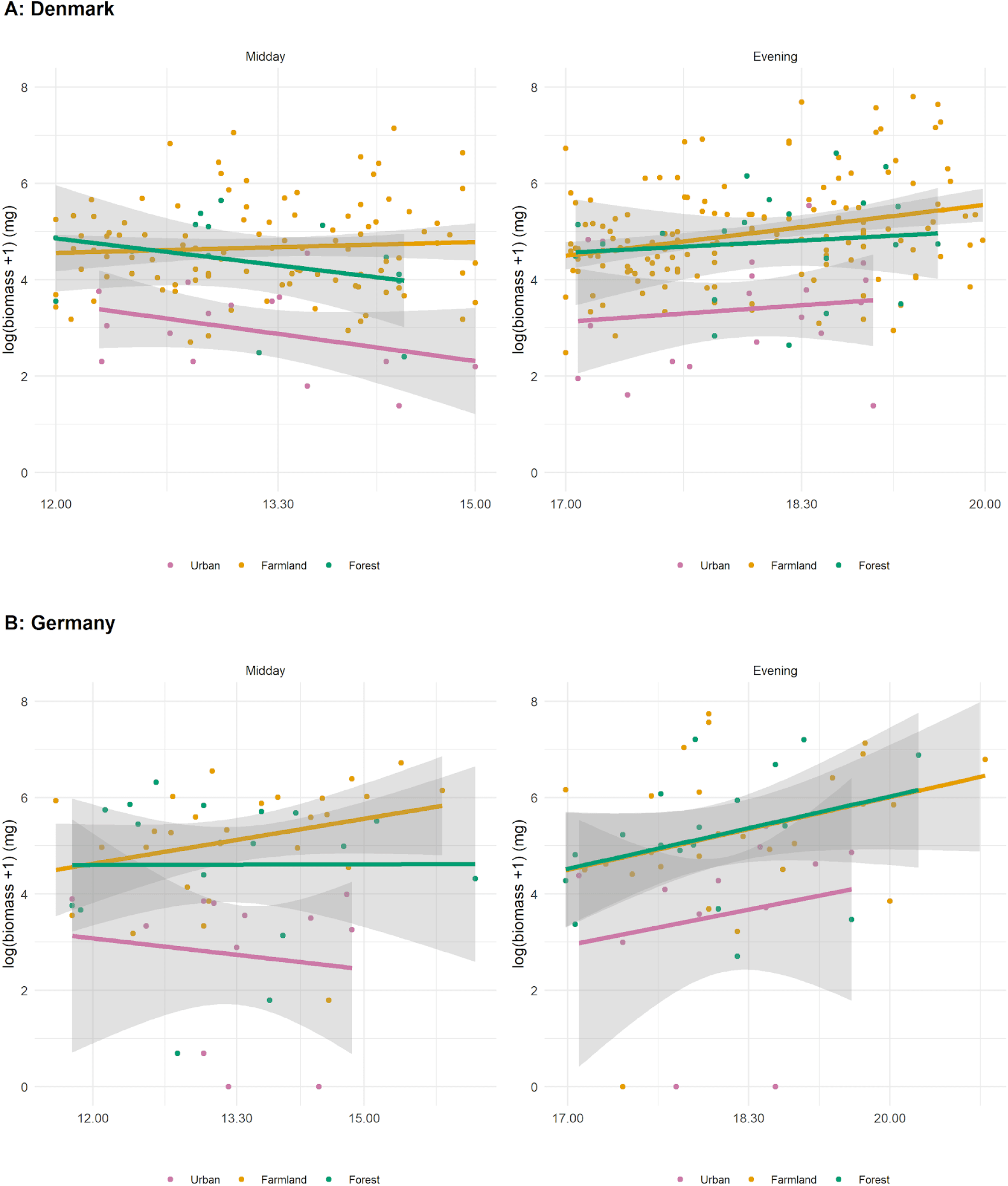
Sampling time effects on insect biomass. A) Denmark and B) Germany: overall effect of sampling time on insect biomass on land covers where the maximum proportional cover could be assigned to a specific land cover category at the 1000 m buffer. Coloured by land covers and shaded areas correspond to the standard error of the fit. We do not show wetland and grassland since these were rarely the dominant land cover along a route.

### Germany

In the best fit model, lower insect biomass was associated with higher urban cover, and higher biomass was found in the evening (Table 1 & Figure 4B). Urban cover and time of day were the only variables retained in the model. The fixed effects of this model explained 30% of the variation in insect biomass. Consistent with these patterns, routes with low biomass samples (within the bottom 20%, <46 mg), were dominated by the urban cover. By contrast, in the routes with high biomass yields (within the top 20%, >502 mg), the mean landscape composition was dominated by farmland cover (see Figure 3C). Similar results were found when land cover variables were square-root transformed, and this reduced multicollinearity problems highlighted by the variance inflation factors found in the full model. Just as for Denmark, there was no evidence of interactions between land cover and time of day, but overall, biomass was higher in the evening compared to midday (Figure 5).

In the full model, including each land cover variable, none of the land cover variables were significant (Table 1). Insect biomass was generally higher in the evening than at midday and further increased with a later start time of sampling during the evening time band (Table 1).

The two main axes identified by the PCA of the land covers were an urbanisation gradient (from urban to farmland) and a forest gradient (from forest to grassland/wetland) (SI II: Figure 2.2B), just as for Denmark. In a model including all control variables, only the urbanisation effect was significant (p=0.036) and not the forest gradient (p=0.20) on insect biomass, again, just as for Denmark. The fixed effects of this model explained 32% of the variation in the data (SI II: Table 2.1).

### Land use intensity in Denmark

The most pronounced effects on insect biomass in both Denmark and Germany were due to urbanisation. To better understand these effects, we considered, within land cover types, a set of sub-types, focused on the intensity of urbanisation. We did the same for sub-types of agricultural land use types. Here, we considered only Denmark for which our sample size was sufficient to allow within land cover type analyses.

When we considered the different subtypes of urban land cover separately, we found a negative effect of urban cover with a high proportional hedge cover and a positive effect of urban areas that had a high cover of commercial areas (Figure 6A & SI III: Table 3.2). However, multicollinearity was an issue for the hedge/urban interaction; hence the result was highly uncertain. For agricultural land use intensity, we found that farmland with a high proportional cover of intensive conventional agriculture had a negative effect on insect biomass; however, multicollinearity was again an issue, and the result was thus highly uncertain. Furthermore, we detected a trend of increased biomass in semi-intensive managed farmland (Figure 6B & SI III: Table 3.4). The partial effect analysis revealed lower insect biomass in intensive conventional agriculture land use compared to the other agricultural land uses.

**Figure 6:**
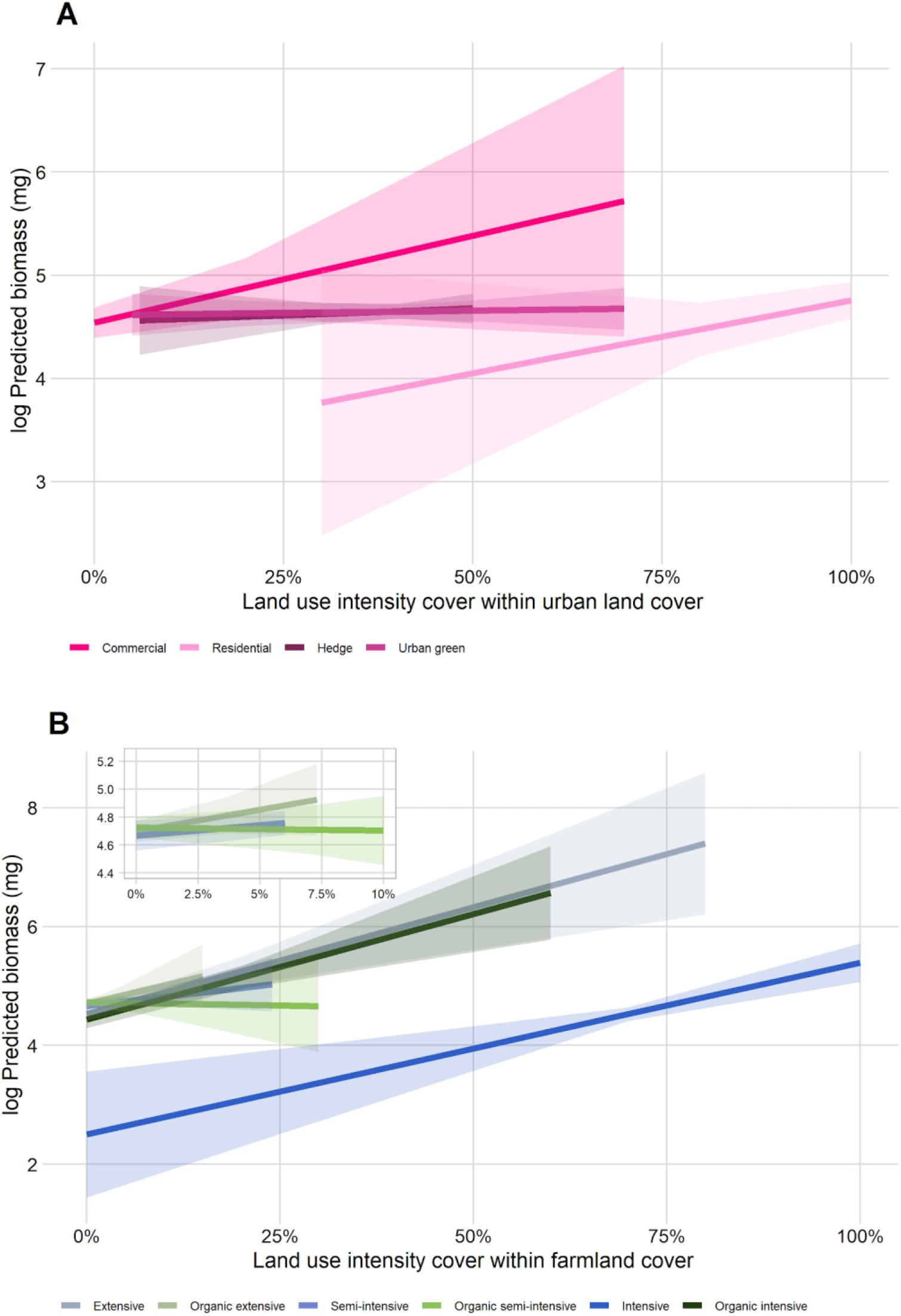
Partial effects of each proportional land use cover in Denmark when all other predictors are held fixed at their means for (A) urban land use within the cover of the 34 most urban routes and (B) farmland land use within the cover of the 255 most agricultural routes, green hues = organic farming, blue hues = conventional farming and general farmland cover. Figure zoomed in for the 10% cover to show the partial effects of the land uses with low coverage on the routes. Only observations in the 1000 m buffer. Predicted log(biomass+1) (mg) on the y-axis and proportional land use cover on x-axis. Shaded areas around each line is the standard error for the fit.

Throughout all models for land cover and land use, the random effects explained between 28-37% of the variation in Denmark (mean site ID variance = 0.07, mean driver ID variance = 0.35) and 47-52% of the variation in Germany (mean site ID variance = 0.44, mean driver ID variance = 0.84).

## Discussion

Using an innovative citizen science method with car nets, we could simultaneously sample over a large geographic area with a total of 278 transects/routes. In doing so, we sampled the flying insects associated adjacent to both public and private lands, including highly populated cities, relatively remote forests and wetlands, and intensive agricultural fields. This sampling approach revealed a consistent spatial pattern in insect biomass across the two countries, namely lower biomass associated with urbanisation.

### Urban land use has a strong negative effect on insect biomass

In both countries, we found the lowest biomass in urban routes compared with all other land covers, confirming our assumption (H2). However, our results did not confirm our assumption of lower biomass associated with increasing land use intensity (H3), since we found a slightly positive effect of commercial areas on insect biomass. The estimated effect of urban areas was still negative after accounting for potential stops during sampling (Figure 4). Our results are consistent with a recent meta-analysis combining data from multiple studies to show an overall negative effect of urbanisation for arthropod diversity and abundance (Fenoglio, Rossetti & Videla, 2020) and the decline of insect diversity with urbanisation at multiple spatial scales (Piano *et al.*, 2020). In large part, this may be due to the reduced biomass and productivity per unit area in urban habitats where much of the landscape is impervious surface, such as cement or rooftops, on which vegetation does not grow. While studies that focus on local, green habitats in cities often find those habitats to be biological diverse (Guénard, Cardinal-De Casas & Dunn, 2015; Turrini & Knop, 2015; Brunbjerg *et al.*, 2018; Mody *et al.*, 2020 Theodorou *et al*., 2020), such studies may risk missing the broader picture, that the unsampled grey spaces of cities are likely to have low biomass, a reality reflected in our results from both Denmark and Germany. Our approach of sampling across a transect of several km, while having limitations, integrates the effects of green and grey spaces on biomass and provides a more complete picture of the mean biomass of insects in a volume of air space over the city. In doing so, it reveals that there is much lower insect biomass in the urban realm than in all other habitats.

### Insect biomass is positively associated with agricultural land cover, but the positive association may be due to specific land use intensities

We found a positive effect of farmland cover on insect biomass in Denmark and a similar tendency was found in Germany, thus not confirming our assumption of lower biomass in agricultural areas (H1). We found this effect, despite a lack of different land use intensity measures available to test, e.g. data on the amount of fertiliser, pesticide application, and pastoral land cover and land use. Although there was some indication that insect biomass generally was lower in intensive conventional agriculture, thus confirming our assumption (H3), the effect was uncertain and perhaps affected by the fact that most of the farmland cover in Denmark is intensive conventional agriculture. Hence, more sampling in agricultural areas might be helpful to test the effect of agricultural management schemes better. Since random effects explained a large part of the variation, e.g. site and sampling variability, more replicates and detailed explanatory variables would benefit future analysis.

Some studies have previously found a positive association between insect biomass and agricultural land use. Hallmann *et al*. (2017) reported substantial declines in insect biomass in protected areas, many of which are cultural habitats in Europe, having been shaped by human activities (Hurford & Schneider, 2007). However, they found weaker declines in areas with a higher proportion of arable land than natural habitats (measured at a 200 m resolution). In a recent global study, van Klink *et al*. (2020) also found weaker declines in terrestrial insect biomass in areas with high crop cover compared to areas with low crop cover at a local scale, but not at a landscape scale.

The relatively high insect biomass found in farmland might be explained by the high availability of food sources for some insects. Indeed, the density of herbivorous insects have been positively correlated with nitrogen loading in the landscape (Haddad *et al.*, 2000; Ritchie, 2000), and nitrogen loading is expected to be highest in areas with high farmland cover. Hence, higher plant biomass, more nutrient input and higher leaf N content may explain the positive correlation of insect biomass with intensive agriculture. Since we focused on biomass, greater biomass might be primarily caused by a few common and highly abundant species, i.e. agricultural pests and their predators. Further work is needed to assess variation in species diversity and composition, which may show contrasting patterns to biomass.

Considering that our car-based sweeping of insects, like most other forms of insect sampling, records activity rather than directly the local abundance of flying insects, an alternative explanation may be that flying insects more easily traverse farmland, while not necessarily breeding or feeding there. To disentangle activity and habitat association, it would be optimal to have additional biomass data from vegetation sweeping along a subset of routes. If our results contrast with patterns derived from other sampling methods, it may suggest that the higher abundance in farmland is rather due to changes in movement behaviour in hostile landscapes.

### Grassland is sparse and an essential habitat for insects

We found higher biomass of insects in forest, wetland and grassland sites in Denmark compared to agricultural sites, similar to a study on Lepidoptera in Britain (Macgregor *et al*., 2019). The grassland land cover category in Denmark consisted of meadows, salt meadows and grassland under the Danish Protection of Nature Act Section 3. Grassland is an important habitat in a European context, with one-third of all grassland used in an agricultural context with management schemes ranging from extensive to intensive land use (Smit, Metzger & Ewert, 2008). Management schemes can have a large impact on insect populations (Plantureux, Peeters & McCracken, 2005). For instance, nutrient loading, i.e. manure or inorganic fertilisers, in managed grasslands, can decrease insect diversity but tends to increase insect biomass and abundance (Haddad *et al*. 2000), so the biomass found in this study could be associated with specific insect groups, e.g. herbivorous and detritivorous species that may thrive under such conditions (Haddad *et al*. 2000).

Management of grasslands in Denmark has changed within the last couple of decades with less grazing by large herbivores leading to lower rates of deposition of organic nutrients, i.e. dung. For example, the dairy cow population grazing outside has decreased by more than a third since the middle of the 1980s (Statistics Denmark, 2019). These management changes have resulted in shifts in nutrient loading amount and frequency. Loss of outdoor grazing dairy cows is associated with a 60% decrease in starling populations (Heldbjerg *et al*., 2016), most likely due to a loss of insects as a food source due to shifts in nutrient loading with consequences for insect diversity and abundance (Plantureux, Peeters & McCracken, 2005). Grasslands in Germany did not differ from other habitats in terms of insect biomass, which may be explained by the difference in the grassland data in Germany compared to Denmark (in Germany, the semi-natural grassland cover could not be distinguished from agricultural grassland, e.g. grass leys). However, it is also possible that this lack of effect was simply an issue of sample size.

### Even a little wetland cover goes a long way

Recent studies suggest that freshwater insects have increased in abundance and biomass over the last decades, possibly due to improved wastewater regulation, such as through the Water Framework Directive (van Klink *et al*., 2020; Termaat *et al*., 2019). However, wetland land cover has decreased by two thirds over the last century in Europe (European Commission, 1995). In Germany, relatively small areas of wetland were sampled by our study, while in contrast, in Denmark, more samples were obtained in proximity to wetlands. In Denmark, despite the low proportional area, wetland had a significant positive effect on flying insect biomass at the local scale, indicating that even small areas of wetland can be important for flying insects, most likely as breeding habitats. In our study, Danish wetland areas had the highest estimated effect on insect biomass compared to the other land covers in the country (Figure 4A).

### A positive effect of forests on insect biomass

We found a positive effect of forest cover on insect biomass in Denmark at an intermediate spatial scale (250 m). In a study of 30 forest sites in Germany, Seibold *et al*. (2019) found complex patterns of insect changes over the last decade. While they found significant overall declines in biomass and species numbers, forest plots exhibited increases in species numbers and abundance of herbivorous species, especially for invasive and potential pest species, as well as for short-range dispersers. In our study, there were no available data on measures of land use intensity in forests; however, especially deadwood volume is expected to have a significant impact on insect biomass, by providing a rich carbon source that is utilised by saproxylic species (Stokland et al. 2012). However, this should be tested by more focused studies incorporating direct measures on the abundance of these habitat types.

### Limitations and opportunities

We found strong trends and effects of land cover types on insect biomass, especially in Denmark. Interestingly, the summer of our surveys was hot and dry. As such, the differences in biomass among the land cover types might have been increased or reduced due to the drought. We found some unexplained site-specific variability (variation between sites and drivers) that may be explained by including temporal effects. As more samples were obtained from Denmark, it was clear, from comparing sample sizes in Denmark and Germany, that increased sample size could also alleviate some of the variations between sites and citizen scientists. Moreover, there inherently are some issues with the independence of hypothesis tests in this study, since the proportional land cover of each land cover was a part of a 100% cover for each route. Thus, an increase in one type of land cover inevitably leads to a loss in others. Hence, both the loss and gain of land cover have to be considered to understand the impact of land use change on insects. This shift is apparent in both countries where increasing farmland cover is associated with decreasing urban cover (SI IV).

### Biomass as a measure of insect community change

We focused our analysis on insect biomass for a number of reasons. Biomass is readily measurable, relates to some ecosystem services (Barnes *et al.*, 2016) and has been reported to be declining in several studies (Hallman *et al.*, 2017, van Klink *et al.*, 2020). Indeed, our findings of reduced biomass in urban areas are consistent with a recent food supplementation experiment suggesting that urban bird populations are more limited by insect food availability than forest bird populations (Seress *et al*., 2020). Moreover, since we found similar effects of land cover for insects flying during midday and evening, there is some evidence that taxa active during different parts of the day are similarly impacted.

However, biomass is only one measure of an insect community and other measures, such as richness and composition, may show contrasting patterns. For instance, biomass may increase, but species richness may decrease if the increase in biomass is driven by common large-bodied or multiple small generalist species. Relationships between body size, rarity and sensitivity to land use will play roles in determining the relationship between biomass and other metrics.

### Car net sample at landscape scales

The car net sampling approach allowed us to sample across a large geographical extent with several citizen scientists sampling under similar conditions in multiple habitats. In addition, car nets provide an alternative to traditional stationary traps, such as Malaise traps or window traps, since they sample at the landscape scale and integrate over local spatial variation. However, the car net shares some of the same sampling bias as other sampling methods, i.e. they sample insect activity, especially taxa that disperse well, rather than the entire insect fauna of the habitat. Moreover, compared to stationary traps, our car net covered quite a short sampling period and specific taxonomic groups like, e.g. butterflies are underrepresented. This is reflected by the biomass of insects which is mostly <5 gram per sample, whereas, e.g. Malaise trap samples may yield up to several hundred grams within the sampling period (Hausmann *et al*., 2020). For this study, the sampling period was usually 10-20 minutes per route; however, the sampling protocol can be designed to have more extended sampling periods with increased frequency, if the purpose is to monitor biomass, abundance and diversity over time. Since we relied on citizen scientists to collect our samples, we designed a sampling protocol that made it possible for as many people as possible to contribute, without specific insect knowledge or expertise. The simple sampling protocol proved to be quite useful, with a response rate, i.e. samples returned to the research institutions, of 86% in Denmark and a response rate of 96% in Germany. The numbers suggest that standardised citizen science schemes can be a powerful approach to monitor insect diversity simultaneously.

## Conclusions

Overall, we found that urbanisation is associated with decreases in insect biomass. Given the rapid growth of cities around the world, this decrease has the potential for widespread consequences and cascading effects on other species. By sampling both grey and green urban areas, we show clear effects of reduced biomass that were not evidenced before. In addition, we show the relative importance of other land covers, particularly in Denmark, where we had more samples. Conventional intensive agriculture tended to be associated with reduced biomass, even when agriculture overall showed relatively high biomass. Because of the difficulty of sampling conventional intensive agricultural fields, we think our results may be the first evidence of such an effect. In Denmark, semi-natural areas tended to have more insect biomass than either urban areas or farmland. Given the geographic extent of urban areas and farmland in Europe, these findings suggests that massive declines in total insect biomass could have already occurred.

## Acknowledgements

We express our sincere gratitude to all volunteers taking part in the insect monitoring program InsectMobile in both countries. We are very grateful for the local nature conservation authorities in Germany who provided sampling permissions in a very short timeframe, and thus made the scoping study possible.

## Author’s contributions

CSS, JHC, RE, JB, CF, APT, AB, RRD conceptualised the project. JCL, CSS, APT, AR, AB, DE VG and SH organised and coordinated the citizen science sampling. JB and VG extracted environmental data for Denmark and Germany, respectively. CSS, LBP and JM carried out the lab work with support from AJH, NMD and TGF. DEB, AB, RRD, APT and CSS developed analysis models and DEB and CSS wrote scripts for statistical analysis and analysed the data. All authors contributed to the development of the manuscript.

## Data accessibility

Files documenting the analyses and all files necessary to reproduce the analyses, including links to raw data and metadata, are available on GitHub (https://github.com/CecSve/InsectMobile_Biomass).

## Appendix A. Supplementary Information

Supplementary data to this article can be found online at:

