## Supplementary Information for "Contrasting impacts of urban and farmland cover on flying insect biomass"

### Supplementary Material

#### I. Land cover and land use intensity definitions

##### Land cover and land use

###### Denmark

All land cover and land use categories in Table 1.1 are compiled to one GIS theme where categories overwrite each other based on the following prioritization:

1) Lake
2) Meadow (excluding grass leys, protected under the Danish Protection of Nature Act Section 3), (Heath - not used for analysis), Bog, Pasture (excluding grass leys, protected under the Danish Protection of Nature Act Section 3), Salt meadow (excluding grass leys, protected under the Danish Protection of Nature Act Section 3)
3) Forest
4) Intensive, Semi-intensive, Extensive (e.g. agricultural fields and grass leys)
5) Build up areas, divided into four categories: Inner city, other areas dominated by multistory buildings (larger cities), areas dominated by low buildings (residential areas), commercial areas
6) Other agricultural areas with no associated crop type

This prioritization means that a lake was categorized as a lake even though it was based within a forest etc.

Ocean cover was removed from land cover assessments based on the GIS layer 'Kommune' (municipality) from The Danish Agency for Data Supply and Efficiency (Danmarks Administrative Geografiske Inddeling (DAGI)). Six small coastal areas were included, however, since the areas were listed as lakes or protected nature reserves in GeoDenmark. The areas in question were mainly Kilen by Struer and Ulvedyb west of Aalborg.

###### Germany

German land data were extracted from the German ATKIS data (2016). See Table 1.1.

Table 1.1 Land cover and land use categories and definitions. The proportion of each category was extracted for four buffer zones (50 m, 250 m, 500 m and 1000 m) for each route.

| Country | Land cover category | Land use category | Category code (if applicable) | Definition (if applicable) | Data source |
|---|---|---|---|---|---|
| **Denmark** | *Urban* | Low building (lav bebyggelse) | FEAT_KODE 9954 | | Geodanmark/FOT |

| | | | | | |
|---|---|---|---|---|---|
| | | Tall building (høj bebyggelse) | FEAT_KODE 9955 | | Geodanmark/FOT |
| | | Commercial (erhverv) | FEAT_KODE 9953 | | Geodanmark/FOT |
| | | Inner city (bykerne) | FEAT_KODE 9952 | | Geodanmark/FOT |
| | *Farmland* | Extensive * | | Grass leys, fallow etc. | Data provided by The Danish Agricultural Agency under the Ministry of Environment and Food of Denmark :<br><br>Field delineations with crop data downloaded from https://kortdata.fvm.dk/download/Markblokke_Marker?page=MarkerGaeldende, theme Marker_2018, date 20180912.<br>Delineation for organic fields downloaded from http://geodata.fvm.dk/geoserver/wfs, theme oekologiske_arealer_2018, date 20191209 |
| | | Organic extensive | | Organic grass leys, organic farms, fallow etc. | |
| | | Semi-intensive * | | berries, orchards, grass leys etc. | |
| | | Organic semi-intensive | | Organic berries, orchards, grass leys etc. | |
| | | Intensive * | | wheat, beans, rape seed, rye etc. | |
| | | Organic intensive | | Organic wheat, beans, rape seed, rye etc. | |
| | *Grassland* | Salt meadow (strandeng) | | | https://danmarksmiljoeportal.zendesk.com/hc/da/articles/360000745778-Arealinformation-Download-af-data BES_NATURTYPER (Beskyttede naturtyper) |
| | | Grasslands under the Danish Protection of Nature Act Section 3 (overdrev) | | | https://danmarksmiljoeportal.zendesk.com/hc/da/articles/360000745778-Arealinformation-Download-af-data BES_NATURTYPER (Beskyttede naturtyper) |
| | | Meadow (eng) | | | https://danmarksmiljoeportal.zendesk.com/hc/da/articles/360000745778-Arealinformation-Download-af-data |

| | | | | | |
|---|---|---|---|---|---|
| | | | | | BES_NATURTYPER (Beskyttede naturtyper) |
| | *Wetland* | Lake (sø) | FEAT_KODE 9942 | | Geodanmark/FOT (FEAT_KODE=9942) & BES_NATURTYPER (Beskyttede naturtyper) Natyp_navn='Sø' accessed from https://danmarksmiljoeportal.zendesk.com/hc/da/articles/360000745778-Arealinformation-Download-af-data |
| | | Bog (mose) | | | https://danmarksmiljoeportal.zendesk.com/hc/da/articles/360000745778-Arealinformation-Download-af-data BES_NATURTYPER (Beskyttede naturtyper) |
| | *Forest* | Forest | | | Geodanmark/FOT Skov (FEAT_KODE=9917) |
| **Germany** | *Urban* | mixed use area | sie_02 | | German ATKIS data (2016) |
| | | specific functional area | sie_02 | | German ATKIS data (2016) |
| | | mining operations | sie_02 | | German ATKIS data (2016) |
| | | residential area | sie_02 | | German ATKIS data (2016) |
| | | industrial/commercial area | sie_02 | | German ATKIS data (2016) |
| | *Farmland* | crops | veg_01 | | German ATKIS data (2016) |
| | | rotational fallow land | veg_01 | | German ATKIS data (2016) |
| | | permanent fallow land | veg_01 | | German ATKIS data (2016) |
| | | land set aside to EU compensation | veg_01 | | German ATKIS data (2016) |
| | | fruit plantage | veg_01 | | German ATKIS data (2016) |
| | | streuobstwiese, i.e., orchards and garden land | veg_01 | | German ATKIS data (2016) |
| | *Grassland* | pasture | 1200 | | German ATKIS data (2016) |
| | | grassland | 1020 | | German ATKIS data (2016) |

| | | | | | |
|---|---|---|---|---|---|
| | *Wetland* | marshes | 43006 | | German ATKIS data (2016) |
| | | water bodies | gew_01_f | | German ATKIS data (2016) |
| | *Forest* | coniferous | 1200 | | German ATKIS data (2016) |
| | | deciduous | 1100 | | German ATKIS data (2016) |
| | | mixed | 1300 | | German ATKIS data (2016) |

*Intensity calculated based on crop code as well as crop names based on the same mapping used yearly for HNV-maps for the Danish Agricultural Agency.

#### Additional land use intensity variables

##### Denmark

Urban green areas (% within each buffer zone for each route) (BU Area - Green ndvix, class 40 & BU Area - Green Urban Atlas, class 41) were extracted from Copernicus ESM (European Settlement Maps 2017 release) in a 2.5 m resolution within GeoDenmark settlement areas. Hedge data (length in meters) was extracted from GeoDenmark for the three categories: living (levende), unknown (ukendt), and not specified (ikke angivet). Aerial images revealed that the majority of the last two categories consisted mainly of living hedge and we found it reasonable to include them. We categorised hedge within settled areas as hedges and outside settled areas as hedgerows  (Table 1.2).

Table 1.2 Extra land use intensity categories for urban included in the land use intensity analysis for urban land use

| **Country** | **Land use intensity variable** | **Description** | **Code** | **Source** |
|---|---|---|---|---|
| **Denmark** | Urban green area | BU Area - Green ndvix & BU Area - Green Urban Atlas | class 40 & class 41 | Copernicus ESM (European Settlement Maps 2017 release) in a 2.5 m resolution (within FOT settlement areas) |
|  | Hedges | length in meters & meters per acre. Living (levende), unknown (ukendt), and not specified (ikke angivet) |  | FOT |

### II. Supplementary figures and models: Land cover analysis

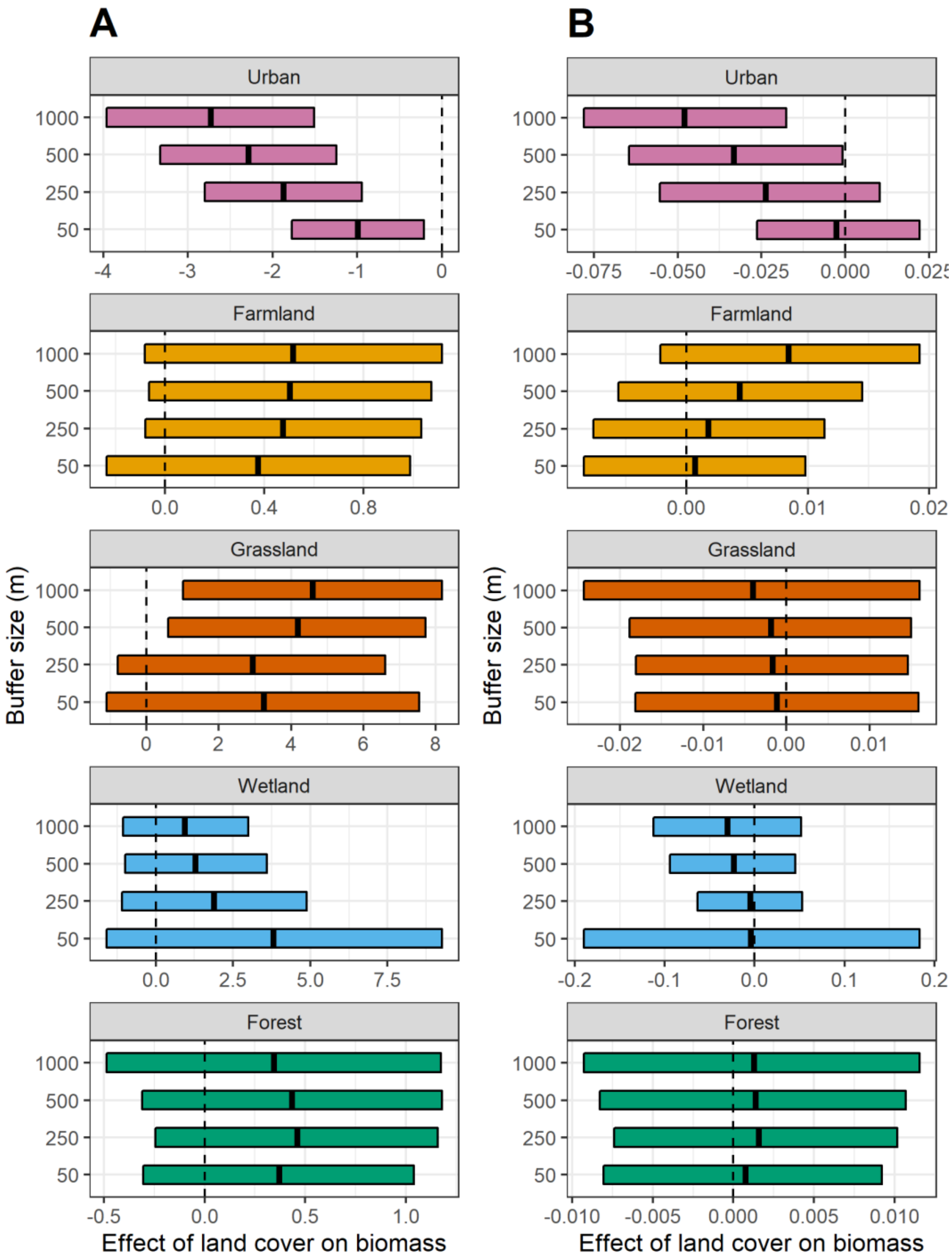


Figure 2.1: Spatial scale effect of different land covers within each buffer zone on insect biomass: A) Denmark, B) Germany. The effect size is the estimated change in log biomass per 1% change in land cover.

PCA analysis for land cover types

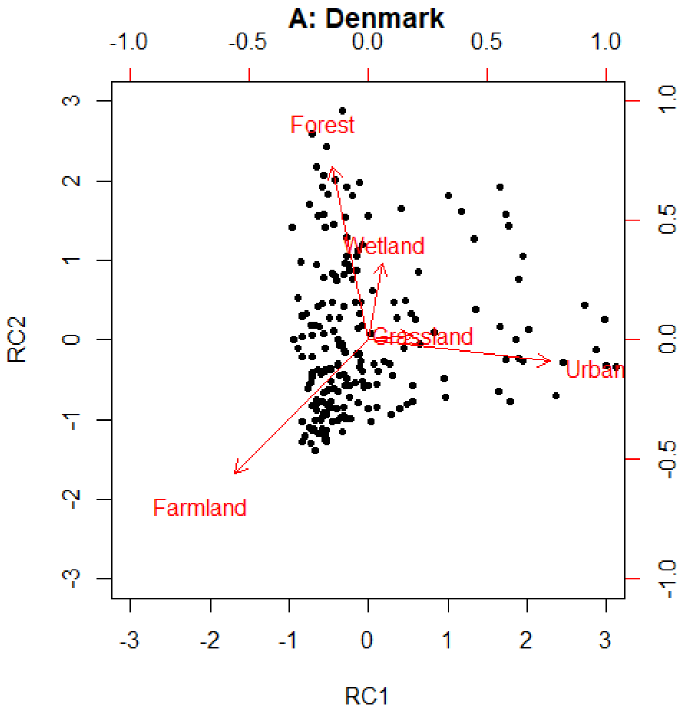


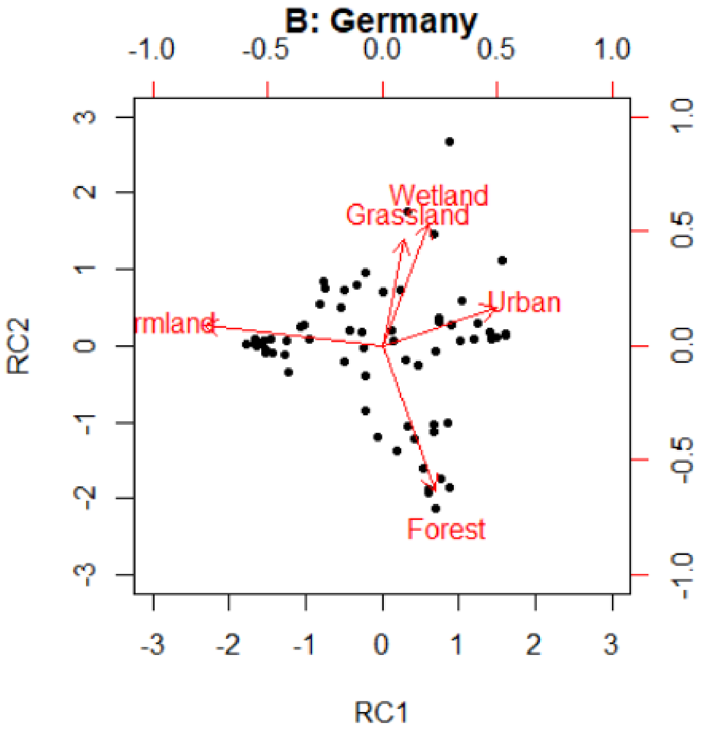

Figure 2.2: Varimax rotated PCA of land covers: A) Denmark, B) Germany

Table 2.1: Regression coefficients of the linear mixed effects model of insect biomass explained by PCA gradient derived land cover. Explanatory variables are axis 1 and 2 from the principal component analysis. All controlling variables were retained regardless of significance. Shown is the mean (standard error) of each regression coefficient.

| **Land cover variable** | **Denmark** (PCA model: 34%) | **Germany** (PCA model: %) |
|---|---|---|
| **Urbanization gradient** | **-0.36 (0.1) *** | **-0.40 (0.19)*** |
| Forest gradient | 0.06 (0.1) | -0.22 (0.17) |
| **Day of month** | **0.03 (0.01) *** | -0.06 (0.04) |
| **Time band**<br>**midday vs evening** | **0.33 (0.1) *** | **0.38 (0.12)*** |
| **Time within band**<br>(change in biomass per minute within time band) | Midday: -0.001 (0.002)<br>**Evening: 0.006 (0.001) *** | Midday: -0.00005(0.0002)<br>**Evening: 0.005 (0.002) *** |
| **Number of Stops** | **-0.39 (0.18) *** | -0.32 (0.20) |

* < 0.05, $^{\Delta}$ < 0.1

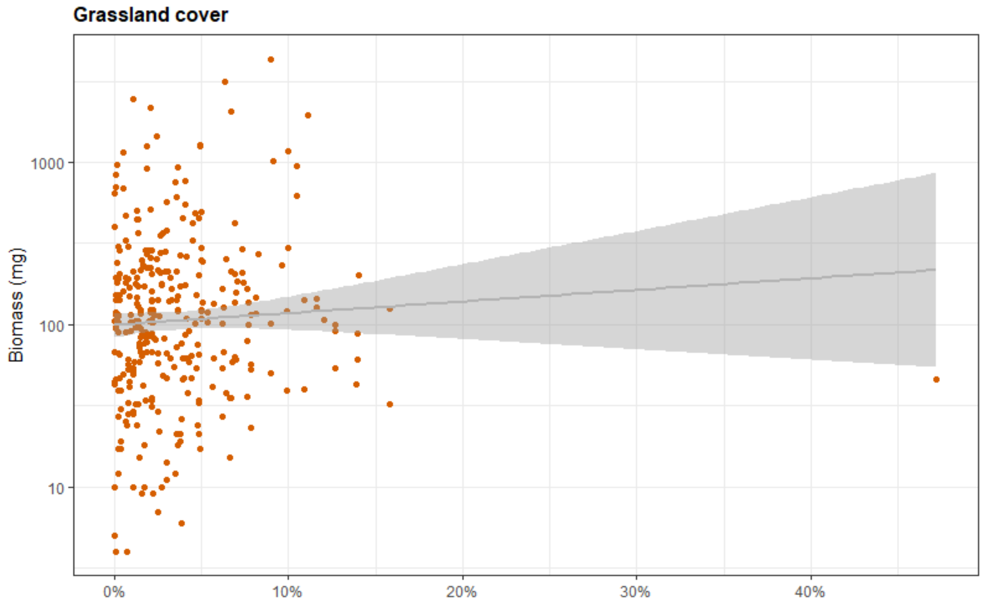


Figure 2.3: Simple regression plot of the Dansih grassland cover with the outlier excluded from the main analysis.

Table 2.3: Model output for the best fit model where the Danish grassland cover outlier is not removed. All land covers are still significant.

| | **Estimate** | **Std. Error** | **df** | **t value** | **P value** | **Significance level** |
|---|---|---|---|---|---|---|
| (Intercept) | 3.41 | 0.49 | 153 | 7.01 | 7.16E-11 | *** |
| Farmland (1000 m) | 1.39 | 0.54 | 150 | 2.57 | 0.011298 | * |

| | | | | | | |
|---|---|---|---|---|---|---|
| Urban (1000 m) | -1.65 | 0.71 | 141 | -2.33 | 0.021378 | * |
| Grassland (1000 m) | 3.37 | 1.49 | 202 | 2.26 | 0.024992 | * |
| Wetland (50 m) | 6.12 | 2.84 | 142 | 2.15 | 0.032992 | * |
| Forest (250 m) | 1.55 | 0.55 | 144 | 2.80 | 0.005869 | ** |
| Time band (evening) | 0.31 | 0.09 | 152 | 3.49 | 0.000641 | *** |
| Sampling day (beginning of June to end of June) | 0.03 | 0.01 | 190 | 4.67 | 5.79E-06 | *** |

Signif. codes: <0.0001 = ***, <0.001 = **, <0.01 = *, <0.1 = .

#### III. Land use intensity analysis

For urban routes, there were strong correlations between urban land cover and the land use intensity variables as well as within some land use intensity variables (urban green space, hedges, and residential areas, and inner city areas and commercial areas) (SI III: Figure 3.1). For farmland, we found a high correlation between intensive conventional agriculture and farmland cover, in accordance with the fact that most farmland in Denmark is intensive agriculture (SI III: Figure 3.5).

#### Urban land use intensity

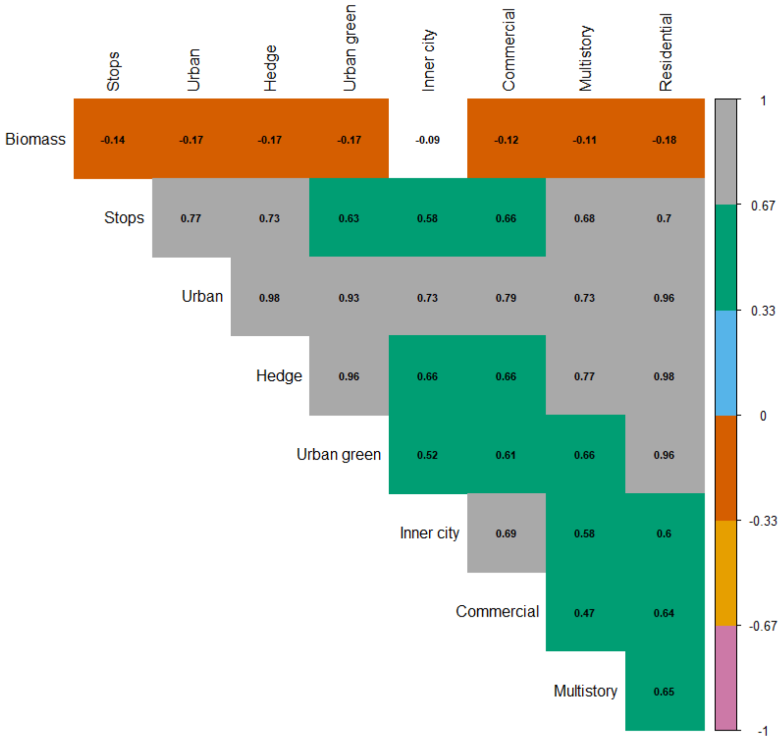


Figure 3.1: Correlation plot of urban land use variables at 1000 m buffer zones (Pearson correlation). High positive correlation = grey. High negative correlation = pink. Significance level <0.05 is indicated by coloured boxes, un-significant boxes are white.

##### PCA

Urban green cover was defined by the first PCA axis in the rotated PCA which was mainly defined by routes with a high proportion of hedges, residential cover and urban green cover. The second axis was mainly defined by inner city cover and commercial cover, both land covers only found in larger cities in Denmark (SI III: Table 3.1 & Figure 3.3). Before PCA rotation, most of the variation for all urban land use variables was defined by the first axis (82%) (SI III: Figure 3.2).

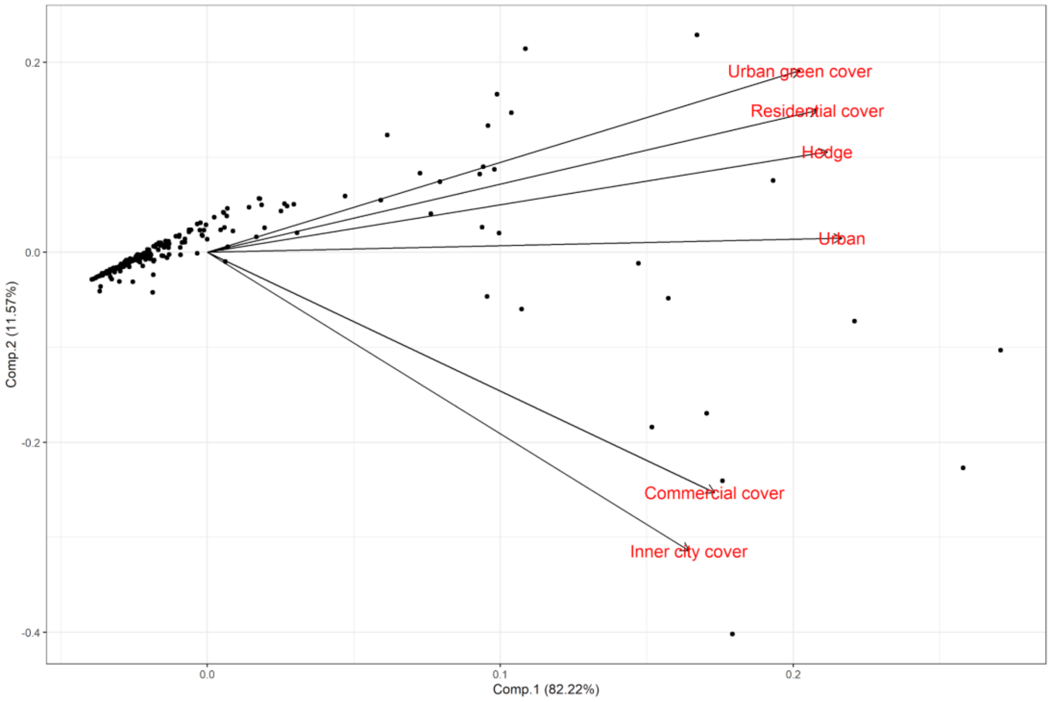


Figure 3.2: PCA of urban land use intensity (prior to PCA rotation).

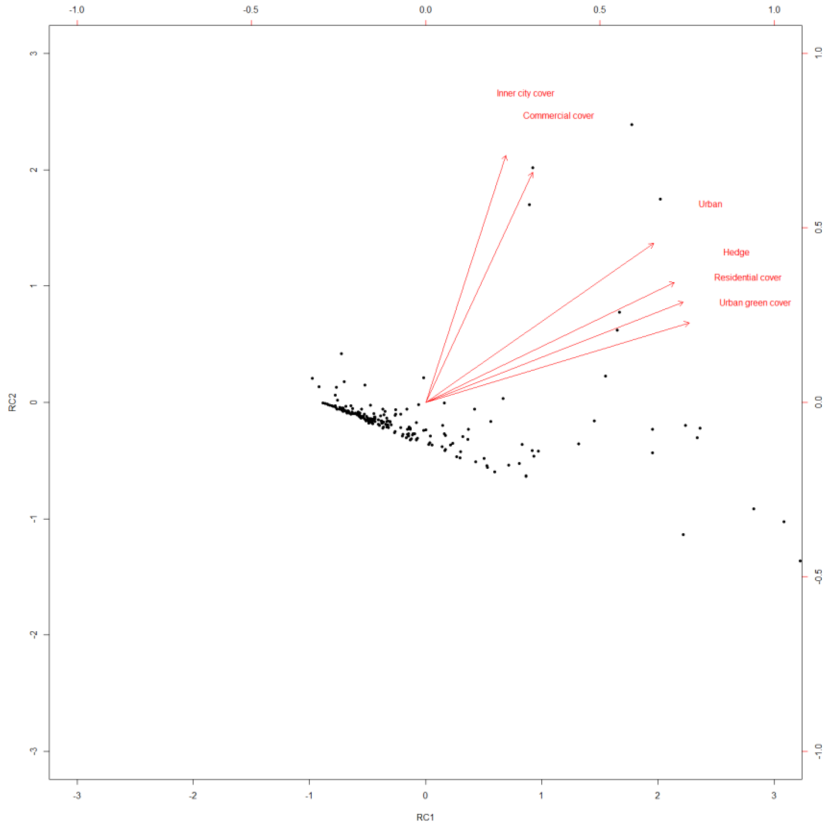


Figure 3.3: varimax rotated PCA of urban land use intensity.

Table 3.1: Principal Component Analysis for urban land use intensity based on rotated PCA.

| Land use cover | RC1 | RC2 | h2 | u2 | com |
|---|---|---|---|---|---|
| Urban | 0.82 | 0.57 | 0.99 | 0.0056 | 1.8 |
| Hedge | 0.89 | 0.43 | 0.98 | 0.0184 | 1.4 |
| Urban green | 0.95 | 0.29 | 0.98 | 0.0247 | 1.2 |
| Inner city | 0.29 | 0.89 | 0.87 | 0.1333 | 1.2 |
| Commercial | 0.38 | 0.82 | 0.83 | 0.1749 | 1.4 |
| Residential | 0.92 | 0.36 | 0.98 | 0.0157 | 1.3 |

#### Correlation & mixed-effects model

Urban land cover was highly correlated with inner city and multistory cover (both only found in larger cities in Denmark, given our study regions), and number of stops (SI III: Figure 3.4).

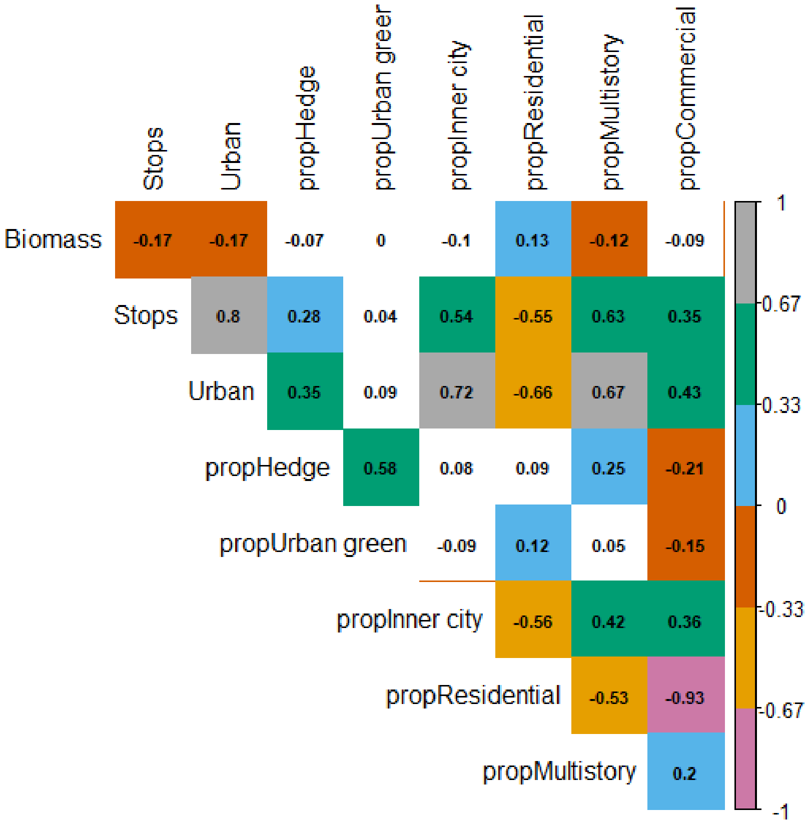


Figure 3.4 Correlation plot of proportional urban land use variables and general urban land cover at 1000 m buffer zones (Pearson correlation). High positive correlation = grey. High negative correlation = pink. Significance level <0.05 is indicated by coloured boxes, un-significant boxes are white. Proportional inner city cover and proportional multistory cover was excluded from the linear-mixed effects model due to high correlation.

In the linear mixed-effects model, we found fixed effects of urban land use and control variables explained 36.4% of the variation. We found a positive effect of sampling date, with more insect biomass in the end of June compared to early June, as well as of time of day, with higher insect biomass in the evening compared to midday, and an increase in biomass the later the evening sampling was carried out (SI III: Table 3.2).

Table 3.2: Regression coefficients of the linear mixed effects model of insect biomass explained by urban land use. Each proportional land use was modelled separately. All controlling variables were retained regardless of significance. Shown is the mean (standard error) of each regression coefficient.

| Model variables | Estimate | Std. Error |
|---|---|---|
| propHedge | 0.001 | 0.001 |

| | | |
|---|---|---|
| propurbGreen | 0.007 | 0.006 |
| propResidential | 0.004 | 0.007 |
| propCommercial | -0.007 | 0.007 |
| **Time band** (midday versus evening) | **0.33** | **0.09 *** |
| Number of stops | -0.29 | 0.21 |
| **Day of month** | **0.03** | **0.01 *** |
| **Urban:propHedge** | **-0.0104** | **4.533 *** |
| Urban:propurbGreen | 0.053 | 0.03 |
| Urban:propResidential | -0..037 | 0.025 |
| **Urban:propCommercial** | **0.065** | **0.028 *** |
| **Time within band (midday)** (change in biomass per minute within time band) | 0.00 | 0.00 |
| **Time within band (evening)** (change in biomass per minute within time band) | **0.01** | **0.00 *** |

* < 0.05, $^{\Delta}$ < 0.1, Generalized variance inflation factors: propHedge/Urban:propHedge = 1.53/28, propurbGreen/Urban:propurbGreen = 1.56/11.6, propResidential/Urban:propResidential = 2.4/19, propCommercial/Urban:propCommercial = 1.88/4.15

#### Farmland land use intensity

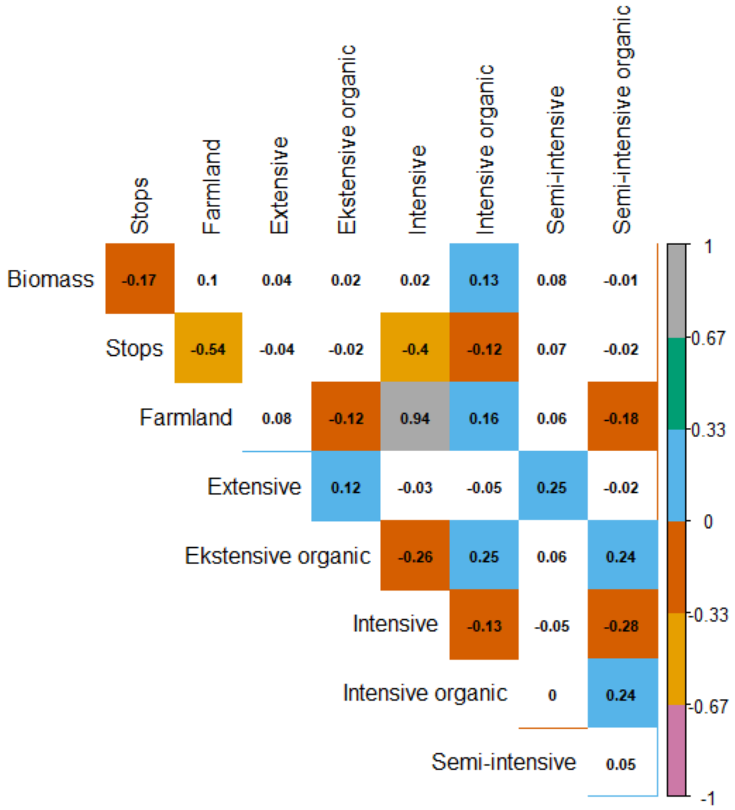


Figure 3.5: Correlation plot of farmland land use intensity variables at 1000 m buffer zones (Pearson correlation). High positive correlation = grey. High negative correlation = pink. Significance level <0.05 is indicated by coloured boxes, un-significant boxes are white.

##### PCA

Before PCA rotation, the farmland/intensive gradient explained 31% of the variation in biomass and the organic/extensive gradient explained 20% of the variation (SI III: Figure 3.6). The rotated PCA appeared similar to the PCA prior to rotation although only loadings for the farmland/intensive gradient were high (SI III Figure 3.7 & Table 3.3).

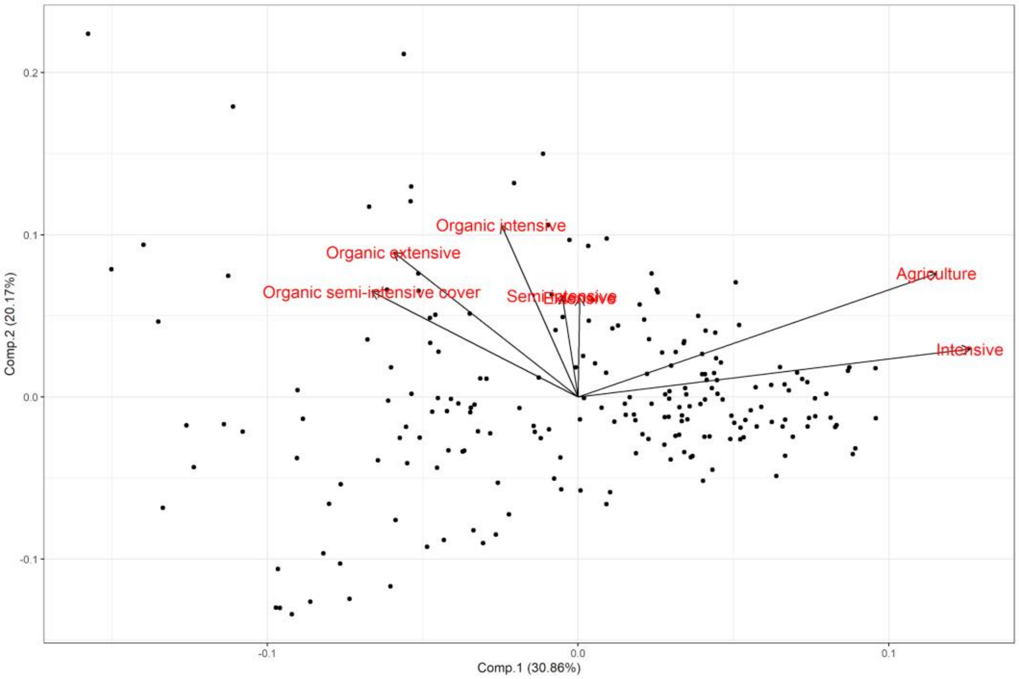


Figure 3.6: PCA of agricultural land use intensity (prior to PCA rotation). Agriculture is the same as general farmland cover used in the farmland cover analysis.

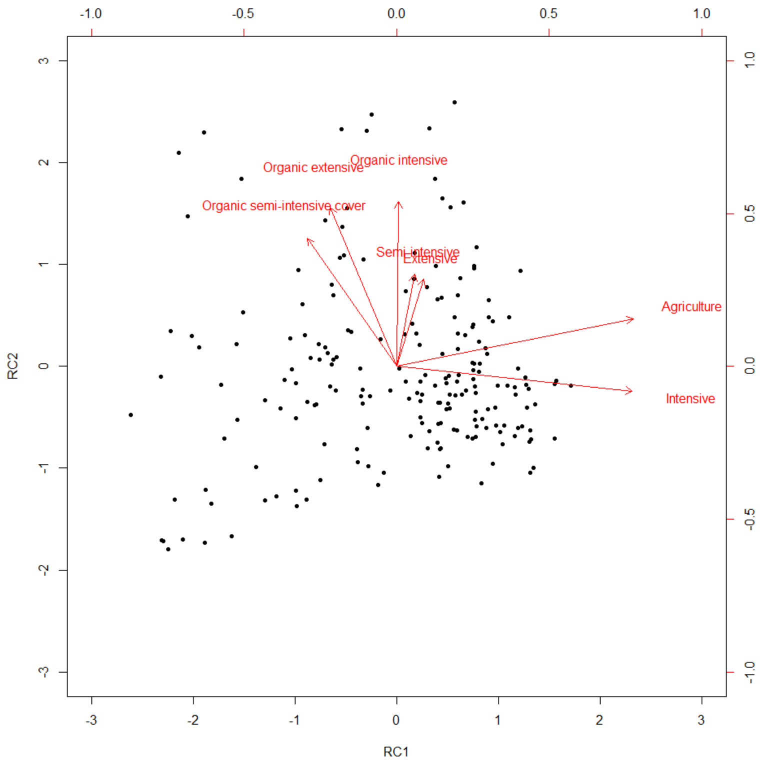


Figure 3.7: Varimax rotated PCA of agricultural land use intensity. Agriculture is the same as general farmland cover used in the farmland cover analysis.

Table 3.3: Principal Component Analysis for agricultural land use intensity based on rotated PCA.

| Land use cover | RC1 | RC2 | h2 | u2 | com |
|---|---|---|---|---|---|
| Farmland (Agriculture) | 0.97 | 0.19 | 0.98 | 0.022 | 1.1 |
| Extensive | 0.11 | 0.36 | 0.14 | 0.860 | 1.2 |
| Organic extensive | -0.27 | 0.65 | 0.50 | 0.501 | 1.3 |
| Intensive | 0.97 | -0.10 | 0.94 | 0.057 | 1.0 |
| Organic intensive | 0.01 | 0.67 | 0.46 | 0.545 | 1.0 |
| Semi-intensive | 0.07 | 0.38 | 0.15 | 0.853 | 1.1 |
| Organic semi-intensive | -0.37 | 0.53 | 0.41 | 0.590 | 1.8 |

#### Correlation & mixed-effects model

None of the land use intensity variables were highly correlated with farmland land cover when we accounted for the proportional cover of the intensity variables within farmland cover. Fixed effects explained 34.8% of the variation. We found a positive effect of sampling date, meaning an increase in insect biomass throughout June, higher insect biomass in evening samples, an increase in insect biomass throughout the evening and a negative effect of potential stops along the routes (SI III: Table 3.4).

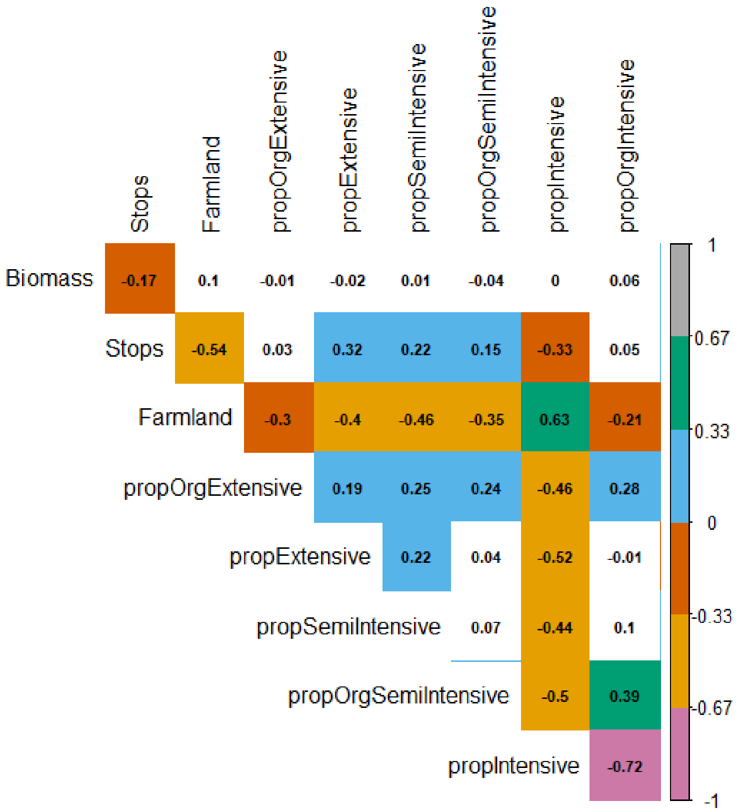


Figure 3.8: Correlation plot of proportional cover farmland land use intensity variables of overall farmland cover at 1000 m buffer zones (Pearson correlation). High positive correlation = grey. High negative correlation = pink. Significance level <0.05 is indicated by coloured boxes, un-significant boxes are white.

Table 3.4: Regression coefficients of the multiple regression model of insect biomass in Denmark. All controlling variables were retained regardless of significance. All land use intensity variables and controlling variables were kept in their original units to facilitate interpretation. Shown is the mean (standard error) of each regression coefficient.

| **Model variables** | **Estimate** | **Std. Error** |
|---|---|---|
| propOrgExtensive | -0.020 | 0.087 |
| propExtensive | 0.004 | 0.011 |
| propSemiIntensive | -0.06 | 0.03 $^{\Delta}$ |
| propOrgSemiIntensive | -0.41 | 0.032 |

| | | |
|---|---|---|
| propIntensive | 0.010 | 0.006 |
| propOrgIntensive | -0.011 | 0.012 |
| **Time band** (midday versus evening) | **0.33** | **0.09 *** |
| **Number of stops** | **-0.60** | **0.17 *** |
| **Day of month** | **0.03** | **0.01 *** |
| Farmland:propOrgExtensive | 0.067 | 0.208 |
| Farmland:propExtensive | 0.026 | 0.036 |
| Farmland:propSemiIntensive | 0.156 | 0.079 $^{\Delta}$ |
| Farmland:propOrgSemiIntensive | 0.127 | 0.137 |
| **Farmland:propIntensive** | **-0.029** | **0.013 *** |
| Farmland:propOrgIntensive | 0.031 | 0.022 |
| Time within band (midday) (change in biomass per minute within time band) | 0.00 | 0.00 |
| **Time within band (evening)** (change in biomass per minute within time band) | **0.01** | **0.00 *** |

* < 0.05, $^{\Delta}$ < 0.1, Generalized variance inflation factors: propOrgExtensive/Farmland:propOrgExtensive = 7/6.6, propExtensive /Farmland:propExtensive = 1.8/1.4, propSemiIntensive/Farmland:propSemiIntensive = 3.7/3, propOrgSemiIntensive/Farmland:propOrgSemiIntensive = 2.14/2, propIntensive/Farmland:propIntensive = 3.6/21, propOrgIntensive/Farmland:propOrgIntensive = 4.4/4.3.

### IV. Stopping effect and land cover correlation

We detected a strong correlation of potential stops and urban sampling at all four buffer zone scales. Furthermore, we find an increasing negative correlation with farmland cover and other land cover categories (Figure 4.1).

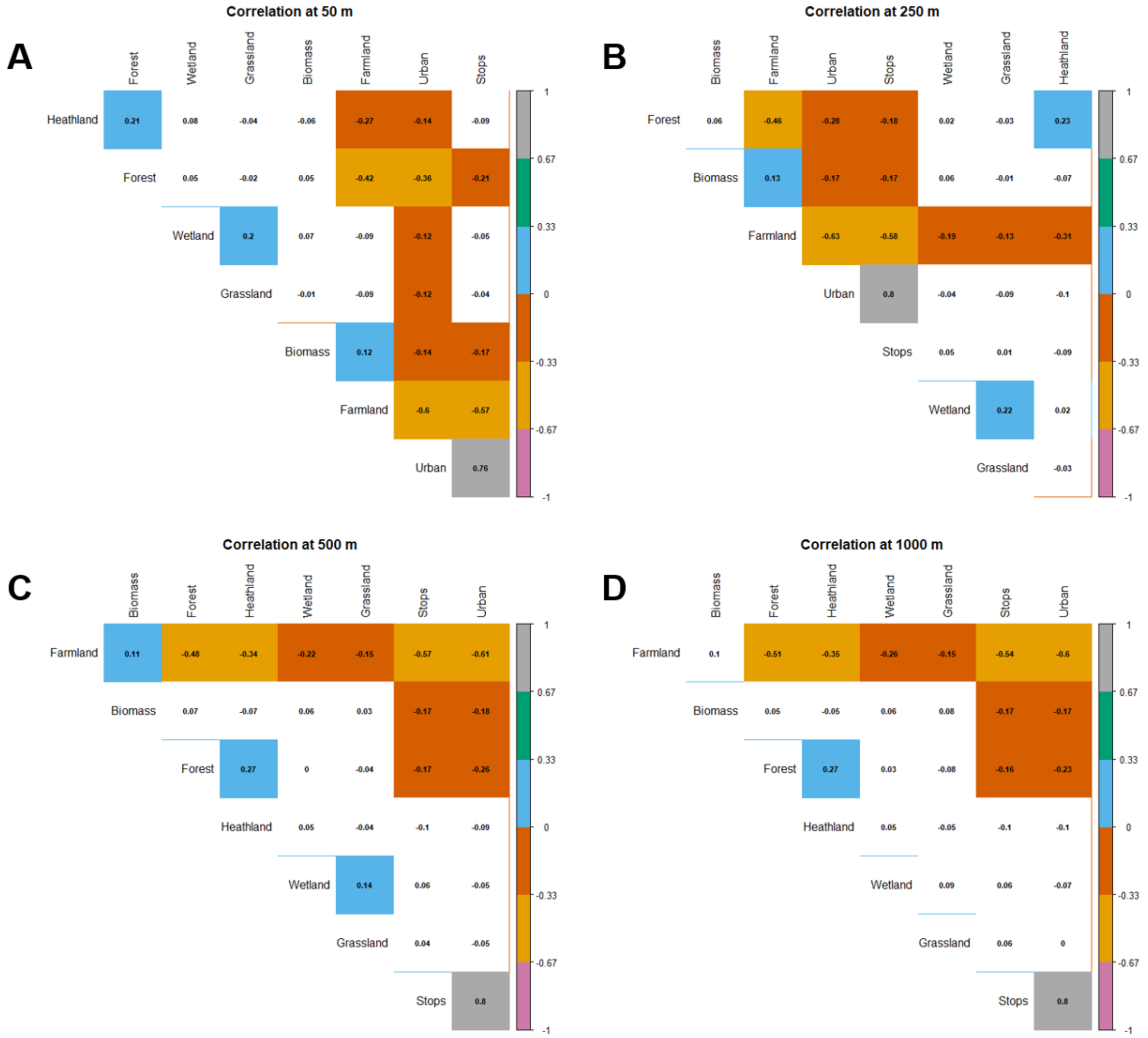
Correlation at 50 m
A
Forest
Wetland
Grassland
Biomass
Farmland
Urban
Stops
Heathland
Forest
Wetland
Grassland
Biomass
Farmland
Urban
Correlation at 250 m
B
Biomass
Farmland
Urban
Stops
Wetland
Grassland
Heathland
Forest
Biomass
Farmland
Urban
Stops
Wetland
Grassland
Correlation at 500 m
C
Biomass
Forest
Heathland
Wetland
Grassland
Stops
Urban
Farmland
Biomass
Forest
Heathland
Wetland
Grassland
Stops
Correlation at 1000 m
D
Biomass
Forest
Heathland
Wetland
Grassland
Stops
Urban
Farmland
Biomass
Forest
Heathland
Wetland
Grassland
Stops
1
0.67
0.33
0
-0.33
-0.67
-1

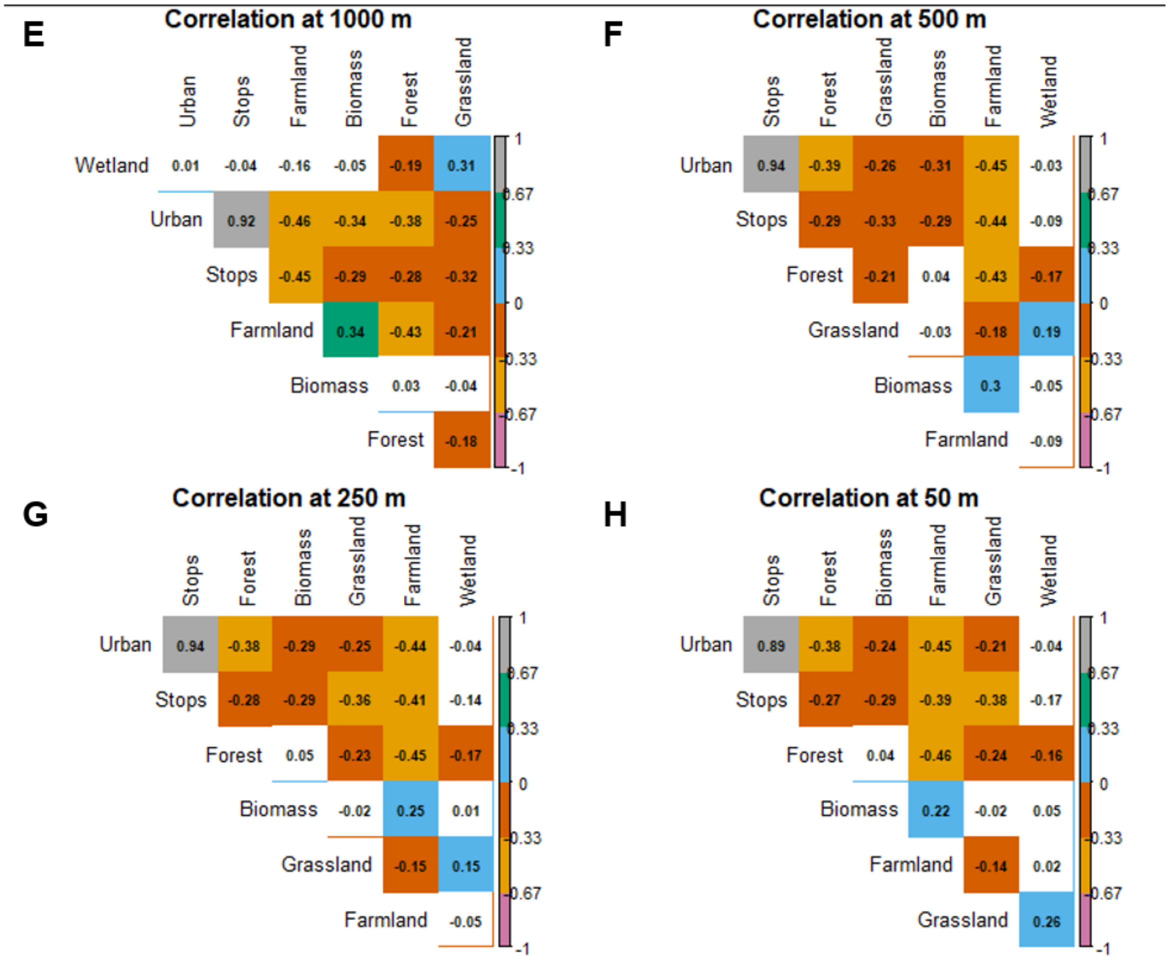


Figure 4.1: Correlation plot of land cover variables, stops, biomass and all four buffer zone sizes (Pearson correlation). A-D) Denmark, E-H) Germany. Significance level <0.05 is indicated by coloured boxes, un-significant boxes are white. High positive correlation = grey. High negative correlation = pink.

### V. Sampling protocol & links to instructional videos

#### Sampling protocol

The following text was part of the sampling protocol sent to the citizen scientists in Denmark, translated from Danish to English:

INSECT MOBILE

CITIZEN SCIENCE
CATCH INSECTS WITH NETS ON THE ROOF OF YOUR CAR
PARTICIPATE AND HELP THE RESEARCHERS

- INSTRUCTIONS -

PACKAGE CONTENTS

We ship two packages with everything you need to collect insects. Check if all the content is in the packages and read the leaflet before you begin to collect insects. Enjoy.

IN THE LARGE PACKAGE YOU WILL FIND

- ❏ tent pole
- ❏ A big net
- ❏ Two sampling bags for Route 1, each containing:
    - ❏ A net-end
    - ❏ A can of alcohol to preserve the insects
    - ❏ Disposable gloves
    - ❏ Data sheet and route description
    - ❏ Pencil to fill in the data sheet
    - ❏ Franked return label to send the packages back
    - ❏ Tape to close the packages

IN THE LITTLE PACKAGE YOU WILL FIND

- ❏ Two sampling bags for Route 2 with the same contents as described above

WELCOME TO THE INSECT MOBILE

Thank you for registering as an Insect Mobile pilot and participating in the insect collection. Together we will investigate flying insects in Denmark, and what habitats they prefer to live in.

Your help is crucial for the Insect Mobile, since it enables us to learn more of the Danish Insect fauna and what efforts we can make to take good care of them.

You collect the insects in the large net, which is mounted on the roof of your car.
In total, you have to drive four trips, and after each trip you have to change small net-end at the end of the large net. The insects that you catch are stored in alcohol and sent to us at the Natural History Museum of Denmark in Copenhagen.

The collection takes place in the month of June and you have to drive on days with good weather (read more on the next page). If it's cold, windy or raining, then the insects do not fly. Once you have driven your trips, send the two packages of insects in spirits, the large net and other equipment you used, back to us. The postage is prepaid.

In this leaflet, you can read more on how to collect insects.

At www.insektmobilen.dk you can watch short instructional videos on how the net is Assembled and mounted, and how to collect an insect sample.

We look forward to seeing what you catch.

Best wishes

Anders, Cecilie and Jonas
Citizen Science

1. CHECK THE CONTENTS OF THE PACKAGES AND READ THE INSTRUCTIONS

When you receive the two packages, make sure that everything is there (see checklist). There are two sampling bags in the large package and two sampling bags in the small package - one for each of your four trips.

Read the points below before you start collecting insects.

2. ROUTES AND WEATHER

We have made two different routes (Route 1 and Route 2) where you have to collect insects. You must drive on each route (1 and 2) and sample insects twice within these times:

- Route 1: Trip 1A between kl. 12-15 and Tour 1B between kl. 17-20
- Route 2: Trip 2A between kl. 12-15 and Tour 2B between kl. 17-20

This means that you have to drive a total of four trips with the net on your car. It is important that you can drive both the early ride (12-15) and the late ride (17-20) on the same day, but you do not have to drive on both route 1 and 2 on the same day. You must drive your trips in the month of June, but only on days when the weather conditions are "good". That means:

- Temperature is 15℃ or more
- No rain
- The average wind speed is equal to or less than 6 m/s

3. HOW TO ASSEMBLE THE NET

Watch instructional video at www.insektmobilen.dk: 'How to assemble the net'

It is important that you do not put the net on your car until you are at the starting point one of your routes.

4. PUT THE NET ON THE ROOF OF YOUR CAR

Watch the instructional video at www.insektmobilen.dk: 'How to mount the net'.
Once you are at the starting point for one of your routes, it's time to place the big net on the roof of your car. Remember to put a new net-end on before you start the collection. Also remember to fill out a data sheet with start time and weather conditions, each time you start collecting insects. Use the enclosed pencil. A ball pen smudges if it comes in contact with alcohol.

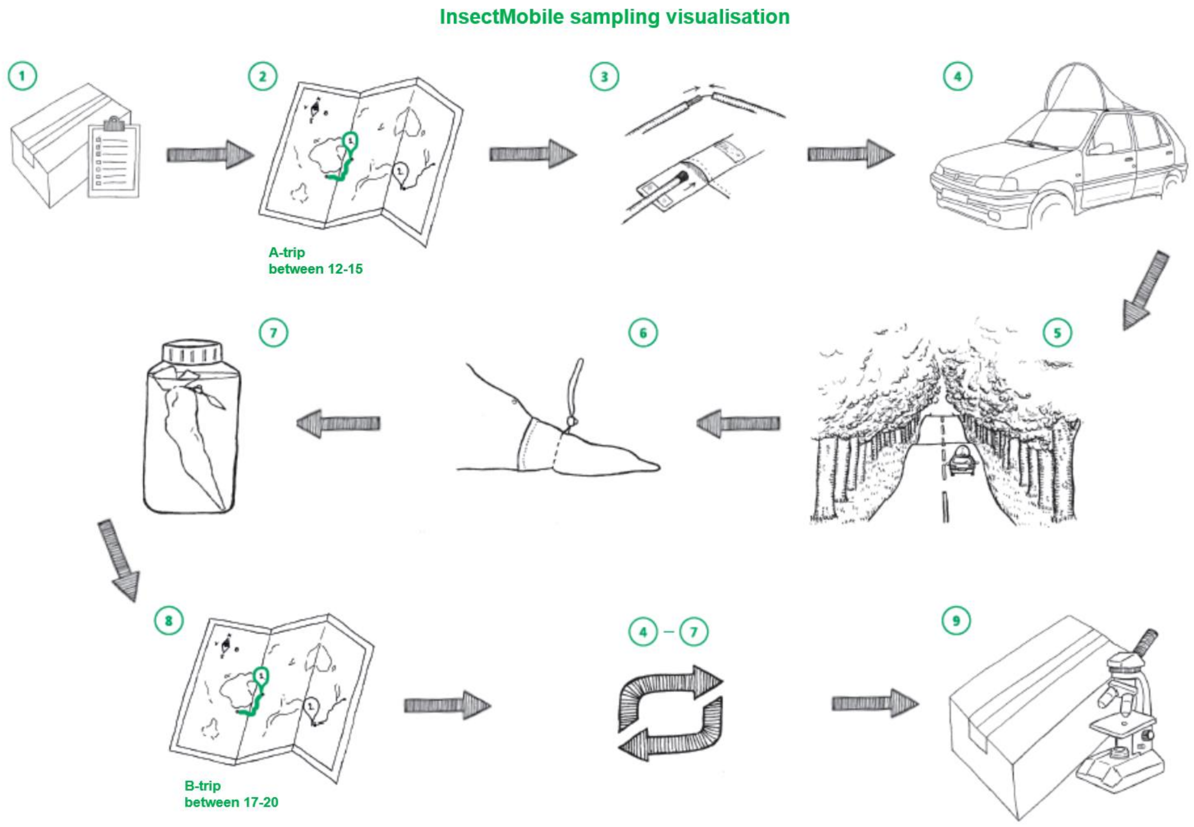


5. HOW TO DRIVE A ROUTE & SAMPLE INSECTS

When the net is on the roof of your car, you may not drive more than 50 km / h.
You sample insects by driving a trip:

- Drive from start point to end point (5 km)
- Turn around at the end point
- Drive back to the starting point (5 km)

Each trip is thus 10 km long. Once you have driven a complete trip, remember to note the end time on the data sheet.

6. TIGHTEN THE NET END SO THAT THE INSECTS DO NOT ESCAPE

Watch instruction video on www.insektmobilen.dk: 'After a sampling trip - securing the sample and sample preparation'

Now the net end must be fastened quickly so that the insects do not escape. Pull the net-end string and tighten well with the string clasp. Then take the net end of the large net.

7. PLACE THE NET-END WITH INSECTS IN ALCOHOL

Unscrew the lid and remove the plastic seal from an alcohol can. Take the net-end with insects and gently lower it into the alcohol can. Pay attention, make sure the liquid does not run over the edge and use rubber gloves if necessary. When the entire net-end is in the can, put the seal back on and screw the lid on tightly.

Congratulations - you have now collected an insect sample with the Insect Mobile!

8. NEXT TRIP

When you are ready to drive your next trip, put a new net-end on the large net and repeat steps 4-7. Remember to fill out a data sheet for each trip you drive.

9. SEND YOUR INSECTS TO US

Once you have run the four trips, pack the large cardboard box with the net, tent poles and two of the cans with insects + data notes. Pack the other two cans with insects + data notes in the small cardboard box. Close both cardboard boxes with tape, put return labels on and drop them off at your local post office or in the parcel box. We have written our address as the receiver  and the postage is paid.

Thanks for the help!

FOLLOW THE INSECT MOBILE

At www.insektmobilen.dk you can follow the project and read more about what happens to the insects when you ship them back to us. You can also follow Insektmobilen on Facebook: www.facebook.com/insektmobilen

And on Instagram where you can share your experiences as an Insect Mobile Pilot with the #insectmobile.

Also stop by: www.facebook.com/statensnaturhistoriskemuseum, where you can read about exciting projects, events and new exhibitions at the Natural History Museum of Denmark.

CONTACT

The project is supported by Aage V. Jensen Nature Foundation

#### Instructional videos

##### How to assemble the car net

https://www.youtube.com/watch?v=z3XNiLukqds&t=0s (in Danish, remember to enable subtitles)

How to mount the car net on a car

https://www.youtube.com/watch?v=eo_81YrpTL4 (in Danish, remember to enable subtitles)

After a sampling trip - securing the sample and sample preparation

https://www.youtube.com/watch?v=JJ-9EZERc6w (in Danish, remember to enable subtitles)
